# Opposing roles of physiological and pathological amyloid-β on synapses in live human brain slice cultures

**DOI:** 10.1101/2024.02.16.580676

**Authors:** Robert I. McGeachan, Soraya Meftah, Lewis W. Taylor, James H. Catterson, Danilo Negro, Jane Tulloch, Jamie L. Rose, Francesco Gobbo, Imran Liaquat, Tara L. Spires-Jones, Sam A. Booker, Paul M. Brennan, Claire S. Durrant

**Author notes:** **Corresponding Author: Dr Claire S. Durrant (****)**. R. McGeachan and S. Meftah contributed equally to this work and are joint first author. Joint first authors have the right to reverse names on the manuscript for CV and presentation purposes.

## Abstract

In Alzheimer’s disease, it is theorised that amyloid beta (Aβ) and tau pathology contribute to synapse loss. However, there is limited information on how endogenous levels of tau and Aβ protein relate to patient characteristics, or how manipulating physiological levels of Aβ impacts synapses, in living adult, human brain. Here, we employed live human brain slice cultures as a translational tool to assess endogenous tau and Aβ release, pathology, and response to experimental manipulation. We found that the levels of Aβ_1-40_ and tau detected in the culture medium depend on donor age, and brain region, respectively. Pharmacologically raising physiological Aβ concentration enhanced levels of synaptic transcripts. Treatment of slices with Aβ-containing Alzheimer’s disease brain extract resulted in postsynaptic Aβ uptake and loss of presynaptic puncta. These data indicate that physiological and pathological Aβ can have opposing effects on synapses in living human brain tissue.

## Introduction

Alzheimer’s disease (AD) is a common neurodegenerative disorder that causes progressive, life-limiting dementia. Neuropathologically, it is characterised by inflammation, atrophy (including loss of synapses and neurons), and accumulation of amyloid-β (Aβ) plaques and neurofibrillary tau tangles^1,2^. Loss of synapses is the best correlate of cognitive decline, and so understanding how early changes to Aβ and tau impact synapse health will be crucial for the development of effective therapeutics^1^. Despite evidence that pathological Aβ and tau can induce synapse dysfunction in model systems^1,3–5^ and reports that novel Aβ-targeting antibodies can, modestly, slow cognitive decline^6–8^, AD research has had limited translational success^1,9,10^. This lack of progress is likely due to many AD models failing to capitulate key aspects of human brain function, including lifespan, brain size, and neuronal diversity^11–14^. We must therefore prioritise exploration of Aβ and tau pathophysiology in the living human brain.

Whilst significant progress has been made in characterising Aβ and tau progression in patients with AD^15^, direct real-time analysis of endogenous Aβ and tau levels, within the healthy human brain presents many challenges. Lumbar cerebrospinal fluid (CSF) displays lowered Aβ_1-42_ and increased phosphorylated tau during normal ageing, which is more pronounced in *APOE* ε4 carriers or AD patients^16–19^. However, CSF measures are an aggregate output from the brain influenced by many factors including: breakdown in the blood-brain barrier^20–22^, circadian rhythms^23–25^, and rates of protein production, degradation or clearance^26–28^. As such, CSF cannot provide direct information on protein levels within brain tissue or the proportion of Aβ or tau arising from different brain regions. By contrast, PET scans, whilst unable to detect soluble forms of protein^5,29–31^, permit spatiotemporal visualisation of fibrillar Aβ and tau burden within the brain. PET scans in ageing adults find that whilst Aβ deposition increases throughout the brain with age, and is worse in *APOE* ε4 carriers^32,33^, tau specifically accumulates within the medial temporal lobe^34,35^. Post-mortem analysis of brain tissue reveals a decline in soluble Aβ levels correlated with increased insoluble Aβ with typical ageing or AD^36–38^. Such findings, however, represent only the end-stage disease and not the dynamics of its progression. Examination of brain interstitial fluid (ISF) provides a much closer representation of protein dynamics in the brain, but these studies are rarely performed due to the invasive nature of ISF sampling restricting analysis to patients with shunts or severe head injury^39,40^. Such limitations mean we still know very little about real-time Aβ and tau dynamics in the living human brain across the lifespan.

Understanding how living synapses respond to real-time changes in Aβ also remains a significant challenge in human research. Studies in rodents^41–44^, non-human primates^45^ and neurons derived from human induced pluripotent stem cells (iPSCs)^46–50^ show that experimentally applied oligomeric Aβ binds to post-synapses and can trigger synaptotoxicity. Whilst analysis of end-stage post-mortem AD brain reveals that oligomeric Aβ accumulates at the synapse^51–53^, direct evidence of synaptotoxicity in human brain is lacking. To our knowledge, the effect of Aβ oligomer application or pharmacologically raising endogenous Aβ concentration has, due to presumed toxicity, never been studied in human patients. The effects of *reducing* Aβ in humans has been restricted to indirect CSF, imaging, or post-mortem studies following clinical trials of Aβ lowering therapies in AD^7,8,54–59^. Interestingly, lowering physiological levels of Aβ in wildtype rodents can disrupt synaptic function, indicating that Aβ may play both physiological and pathological roles at the synapse^3^. Understanding how synapses in live human brain respond to both pathological and physiological alterations to Aβ will fill key knowledge gaps that could refine therapeutic design for AD.

In this study, we harness live human brain slice cultures (HBSCs)^60–63^ to explore how endogenous Aβ and tau release from brain tissue is impacted by patient age, sex, brain region, and *APOE* genotype. We establish HBSCs as a translational drug screening tool, demonstrating that endogenous Aβ release can be bi-directionally manipulated by pharmacological compounds, impacting levels of synaptic transcripts. Using array tomography microscopy, we demonstrate that, within 72 hours, AD-derived Aβ binds to post-synaptic compartments in live human brain tissue and results in loss of pre-synaptic puncta. Finally, we find evidence of Aβ plaques and tau tangles in a subset of our samples, with levels of pathology in the slice correlating with Aβ_1-42_/Aβ_1-40_ ratio in the culture medium. Our data underscores the potential of HBSCs to investigate unexplored aspects of human pre-clinical AD pathology.

## Results

### Preservation of cellular diversity and function in HBSCs at 7 days *in vitro*

Over the course of this study, we generated slice cultures from surplus neocortical tissue removed to obtain access to the tumour for debulking surgery of glioblastoma (16/29), glioma (9/29) or brain metastases (4/29), from 29 patients (**Fig. 1a**). Our total cohort comprised of 10 females and 19 males with an average age of 57 years old (ages ranging from 28-77 years old) (**Table 1**). The majority of samples were from temporal (10/29) and frontal (11/29) brain regions, with a subset of samples derived from parietal or occipital lobes (8/29). First, we established whether key cell types in the brain were preserved in HBSCs for at least 7 days in vitro (*div*). MAP2 and NeuN positive neurons were preserved in HBSCs, across the cortical layers (**Fig. 1b-d**). We also detected microglia in different states, using the classical pan-microglia marker Iba1^64^ (**Fig.1e**) alongside the pyrogenic receptor P2RY12 (known to be expressed on the ramified processes of microglia^65^) (**Fig. 1f**). GFAP-positive astrocytes were present throughout the slice tissue, and could be observed in contact with blood vessels (**Fig. 1g**). At 7 *div* we were able to detect spontaneous synaptic activity and evoke action potentials using whole-cell patch clamp recordings of individual neurons (**Fig. 1h,i**).

**Figure 1:**
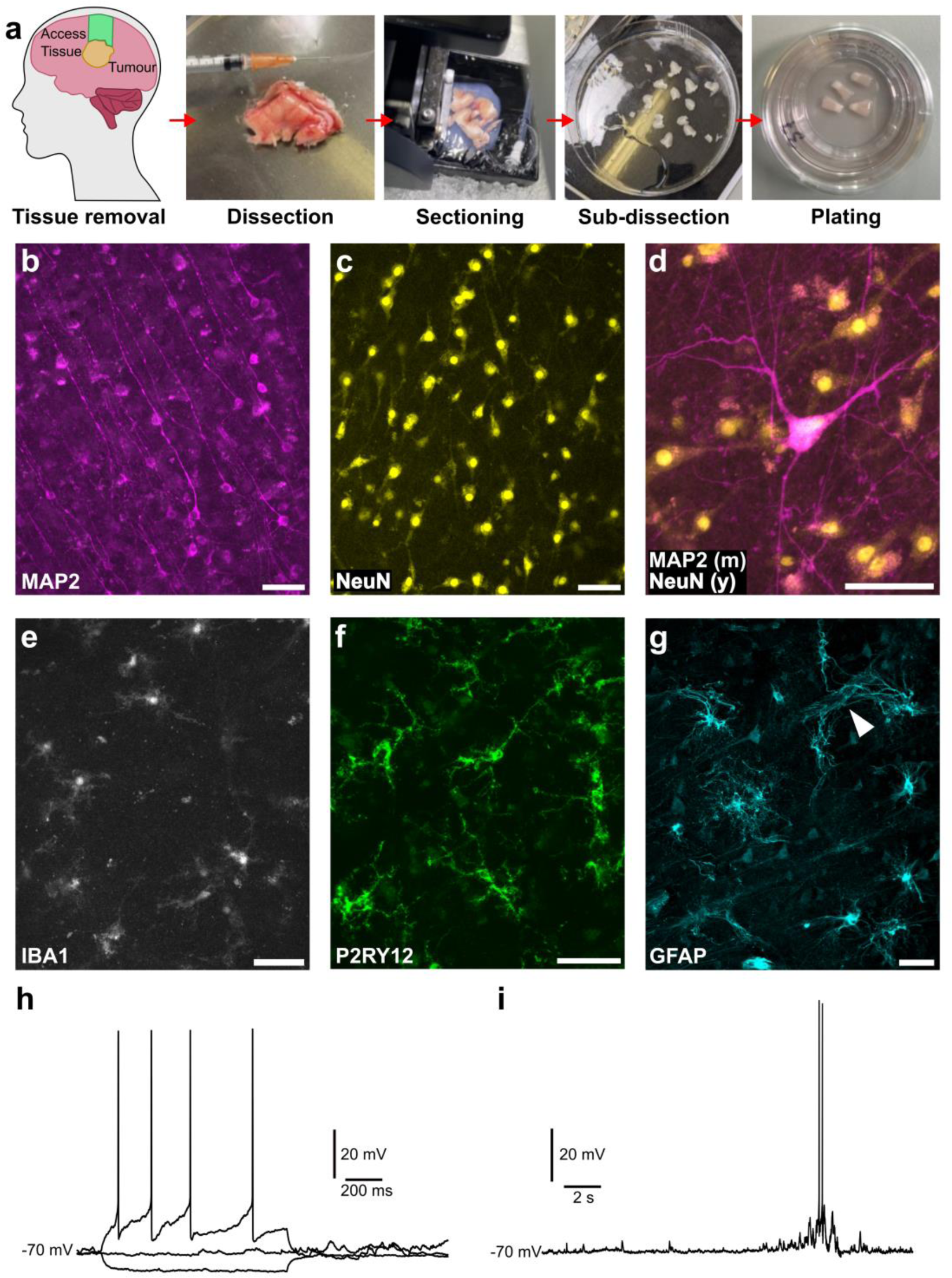
Characterisation of human brain slice cultures. **(a)** An overview of human brain slice culture generation, starting from access tissue removal during brain tumour surgery through to slice culture plating in dishes. (**b-g**) 7 div immunofluorescence images of MAP2 **(b)**, NeuN **(c)**, MAP2 (magenta) and NeuN (yellow) merged **(d)**, IBA1 **(e)**, P2RY12 **(f)** and GFAP **(g)**. A white arrowhead highlights astrocyte end feet wrapping around a putative blood vessel **(g)**. Scale bars are 50 µm in length shown at bottom right corner of image. **(h,i)** Example electrophysiological properties of slice cultures at 7 div. **(h)** Responses to hyperpolarising and depolarising current injections, eliciting action potentials at depolarising currents. **(i)** Spontaneous synaptic properties at rest showing small and large coordinated levels of synaptic drive.

**Table 1:**
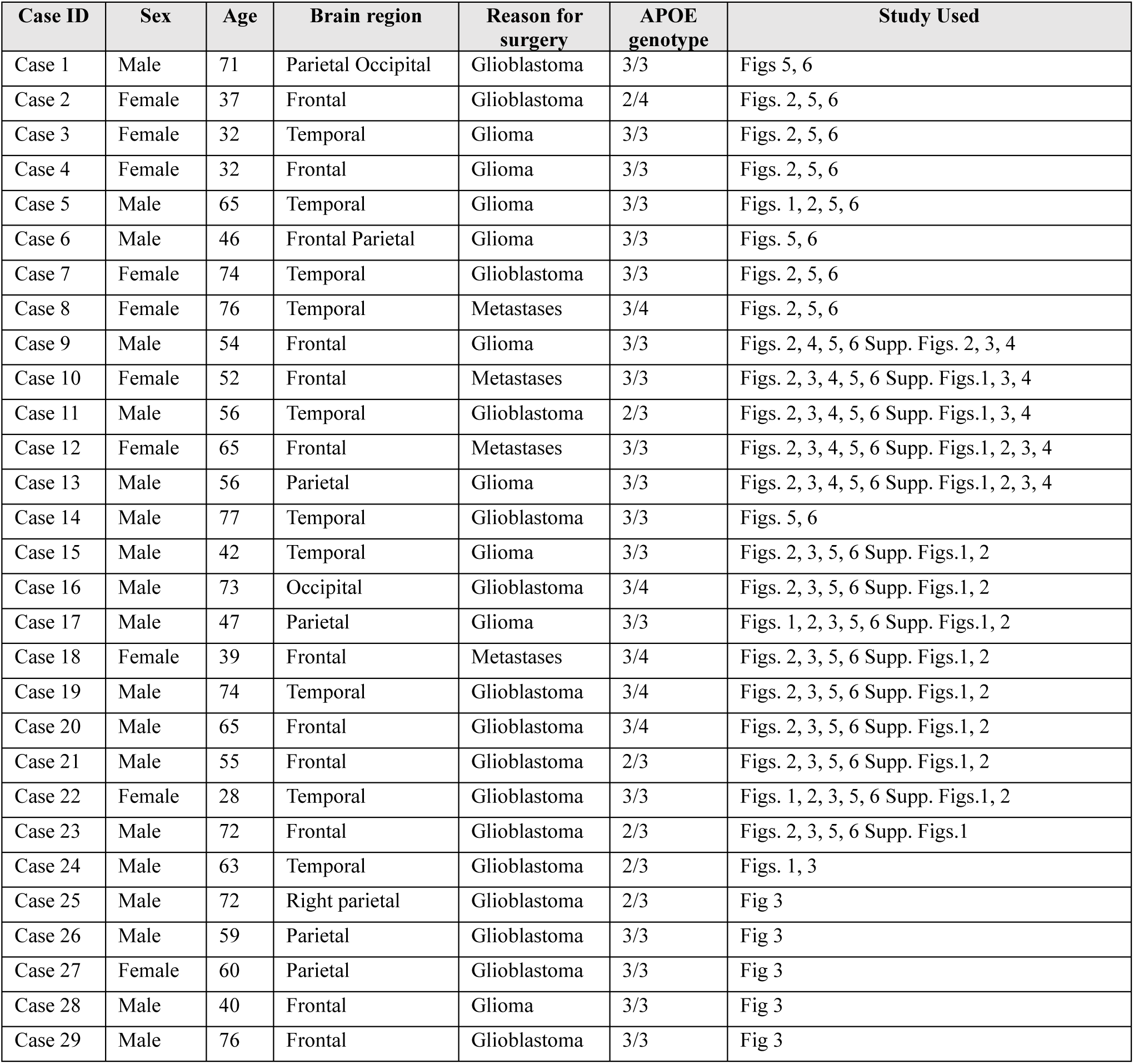
Case details for human brain slice cultures.

### The impact of patient characteristics on basal release of Aβ and tau

Next, we took advantage of the biological diversity of our HBSCs, to assess whether differences in brain region, donor age, sex, or *APOE* genotype impact the basal release of key AD-associated proteins. Tissue from 21 different cases, ages ranging from 28-77, were cultured for 7 *div*, following which the culture medium was collected for analysis of Aβ_1-40_, Aβ_1-42,_ and total tau by ELISA. Endogenous release of these proteins, normalised to total protein, was readily detected in culture medium. We found a trend for a decline in Aβ_1-40_ detected in the culture medium with increasing patient age (**Fig. 2a**), whilst the observed levels of Aβ_1-42_ remained stable with age (**Fig. 2b**). The levels of Aβ_1-40_ were around 16 times higher than the levels of Aβ_1-42_ at baseline, with an average concentration of 16.6+/-5.8 pM and 1.1+/-1.9 pM respectively. Aβ_1-42_/Aβ_1-40_ ratio remained relatively stable with donor age (**Fig. 2c**). We found no correlation between patient age and the concentration of total tau protein in the medium, with an average detected concentration of 1.4+/-0.9 nM (**Fig. 2d**).

The majority of our samples for this experiment were taken from either temporal lobe (9/21) or frontal lobe (9/21), with only 3 samples taken from parietal or occipital regions (classed as “other” in this study). We found no effect of brain region on the levels of Aβ_1-40_ (**Fig. 2e**) or Aβ_1-42_ (**Fig. 2f**) detected in the culture medium, and no impact on the Aβ_1-42_/Aβ_1-40_ ratio (**Fig. 2g**). We found a significant effect of brain region for tau, with temporal lobe samples having higher levels of tau in the culture medium, when compared to samples originating from the frontal lobe (**Fig. 2h**). This is especially interesting in light of the fact that tau pathology in AD generally originates in the medial temporal lobe^66–68^

*APOE* polymorphic alleles are the main genetic determinant of late-onset AD risk, with individuals carrying the ε4 allele at greater risk of developing the disease, and those with ε2 having lower risk^69,70^. *APOE* genotype distribution for our samples was representative of what we would expect to see in a white British population, with ε3/ε3 being the most common genotype^70^. ε2/ε2 or ε4/ε4 genotypes were not represented in our donor samples. We found no overt impact of *APOE* genotype on the levels of Aβ_1-40,_ Aβ_1-42_ or total tau detected in the culture medium in our samples (**Fig. 2i-l**).

**Figure 2:**
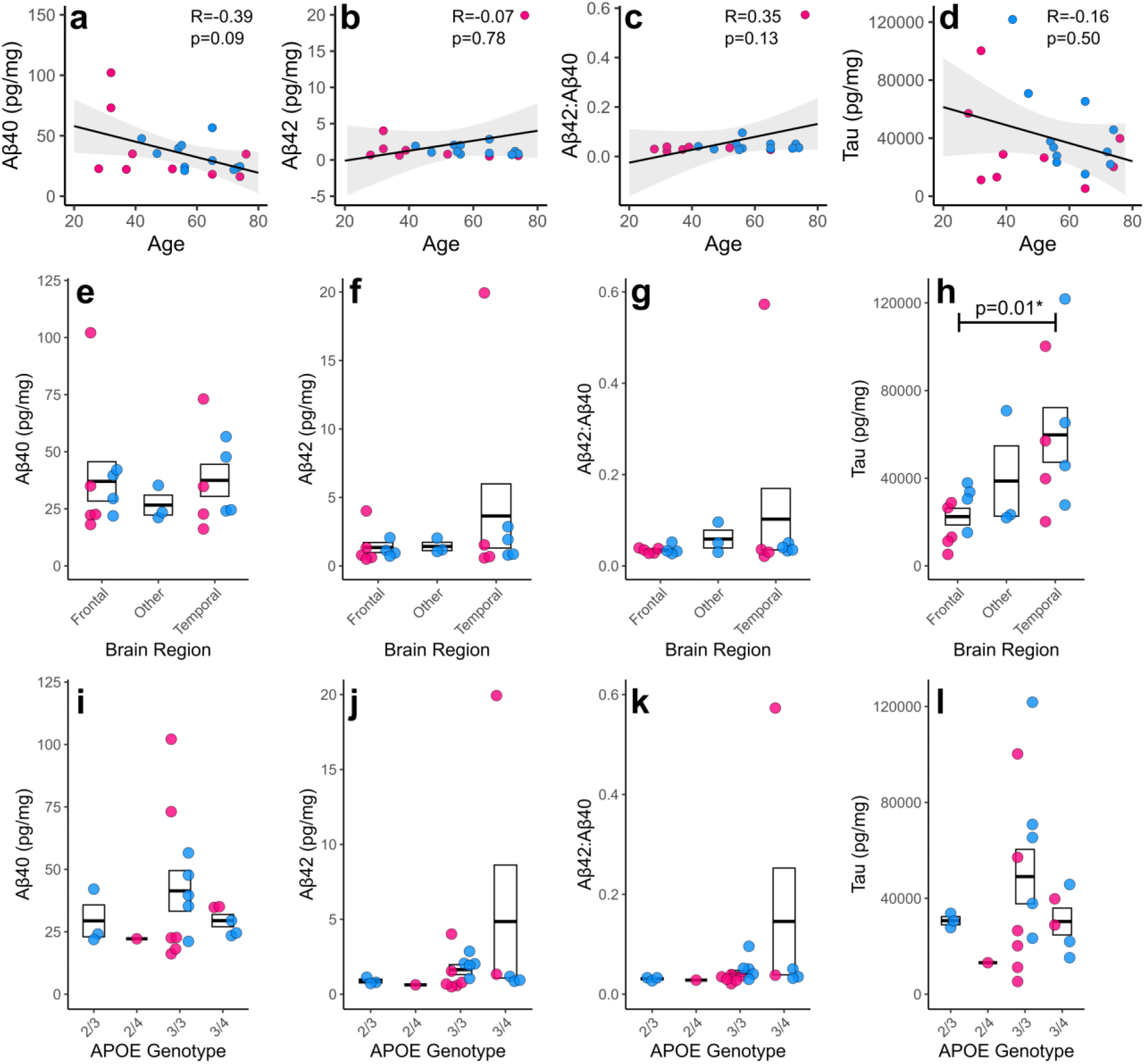
Relationship between Aβ, tau, and patient characteristics. Proteins of interest in the culture medium were measured by ELISA and normalised to total protein content in the medium. **(a-d)** Scatter plots showing the correlations between age of patient and Aβ1-40 (rs(18)=-0.39, p=0.09) **(a)**, Aβ1-42 (rs(18)=-0.07, p=0.78) **(b)**, Aβ1-42/Aβ1-40 ratio (rs(18)=0.35, p=0.13) **(c)**, and total tau (rs(18)=-0.16,p=0.50) **(d)** detected in the culture medium. Statistics: Spearman’s rank correlation due to non-normal distribution. **(e-h)** Box and dot plots assessing the relationship between brain region and Aβ1-40 (log, χ²(2,20)=1.49, p=0.47) **(e)**, Aβ1-42 (log, χ²(2,20)=0.86, p=0.64) **(f)**, Aβ1-42/Aβ1-40 ratio (gaussian, χ²(2,20)=1.26, p=0.53) **(g)**, and total tau (log, χ²(2,20)=17.04, p<0.005***) **(h)** in the culture medium. Statistics: GLMM with format var ∼ Brain region + Sex + (1|Age). **(i-l)** Box and dot plots assessing the relationship between APOE genotype and Aβ1-40 **(i)**, Aβ1-42 **(j)**, Aβ1-42:1-40 ratio **(k)**, and tau **(l)** detected in the culture medium. Pink dots are for female cases and blue dots for male cases. Box represents standard error of the mean, with thick lines representing the mean.

### Aβ release can be bi-directionally manipulated and alters mRNA expression of synaptic genes

Having determined that the endogenous release of AD-relevant proteins could be readily detected in our sample set, we next sought to explore whether we could pharmacologically alter Aβ in HBSCs. Previous studies in mice or primary cultures have shown that inhibition of the beta-secretase beta-site APP cleaving enzyme (BACE1) significantly reduces Aβ production^58^, whilst inhibition of metalloproteases reduces Aβ breakdown^71–75^. Here, we employed a repeated measures design such that tissue from the same patient was split into three independent HBSC dishes. HBSCs from the same patient were then treated with either 5 μM of the BACE1 inhibitor LY2886721, 100 μM of the metalloprotease inhibitor Phosphoramidon, or an untreated medium-only control. Culture medium was collected after 7 *div*, with the measured levels of the protein of interest normalised to the corresponding medium only control. Secreted, soluble levels of Aβ_1-40_ (**Fig. 3a**) and Aβ_1-42_ (**Fig. 3b**) detected in the culture medium were significantly increased by Phosphoramidon treatment and reduced by BACE1 inhibition. The Aβ_1-42_/Aβ_1-40_ ratio remained unchanged regardless of treatment (**Fig. 3c**), suggesting the two isoforms were proportionally impacted. The concentration of tau detected in the culture medium was unchanged by treatment (**Fig. 3d**). Absolute data (not normalised to medium-only control) is available in (**Supp. Fig 1**). The levels of APP protein, as measured by Western blot, remained stable with treatment (**Supp. Fig. 2**), confirming that BACE1 or metalloprotease inhibition acts downstream of APP to alter Aβ levels. Guanidine extraction of insoluble Aβ from the slice tissue showed no impact of Phosphoramidon or BACE1 treatment on the detected levels of Aβ_1-40_ or Aβ_1-42_ (**Fig 3e**). There was, however, a significant treatment effect on the percentage area of Aβ within the slice when measured by MOAB-2 (pan-Aβ antibody that does not detect APP) immunostaining (**Fig. 3f**). Interestingly, Phosphoramidon treatment resulted in increased expression of the synaptic transcripts synaptophysin (*SYP*) and synaptotagmin 1 (*SYT1*), with *SNAP25* also having a significant effect of treatment (**Fig. 3g**). There was also a significant treatment effect on *IBA1* mRNA expression. The levels of *GAD2*, *GFAP* and *C1QA* mRNA remained unchanged by treatment (**Fig. 3g**). Western blots revealed that neither BACE1 inhibitor nor Phosphoramidon treatment impacted levels of neuronal proteins Tuj1 or PGP9.5, synaptic proteins synaptophysin and PSD95, or astrocytic proteins GFAP or YKL-40 (**Supp. Fig. 2**).

**Figure 3:**
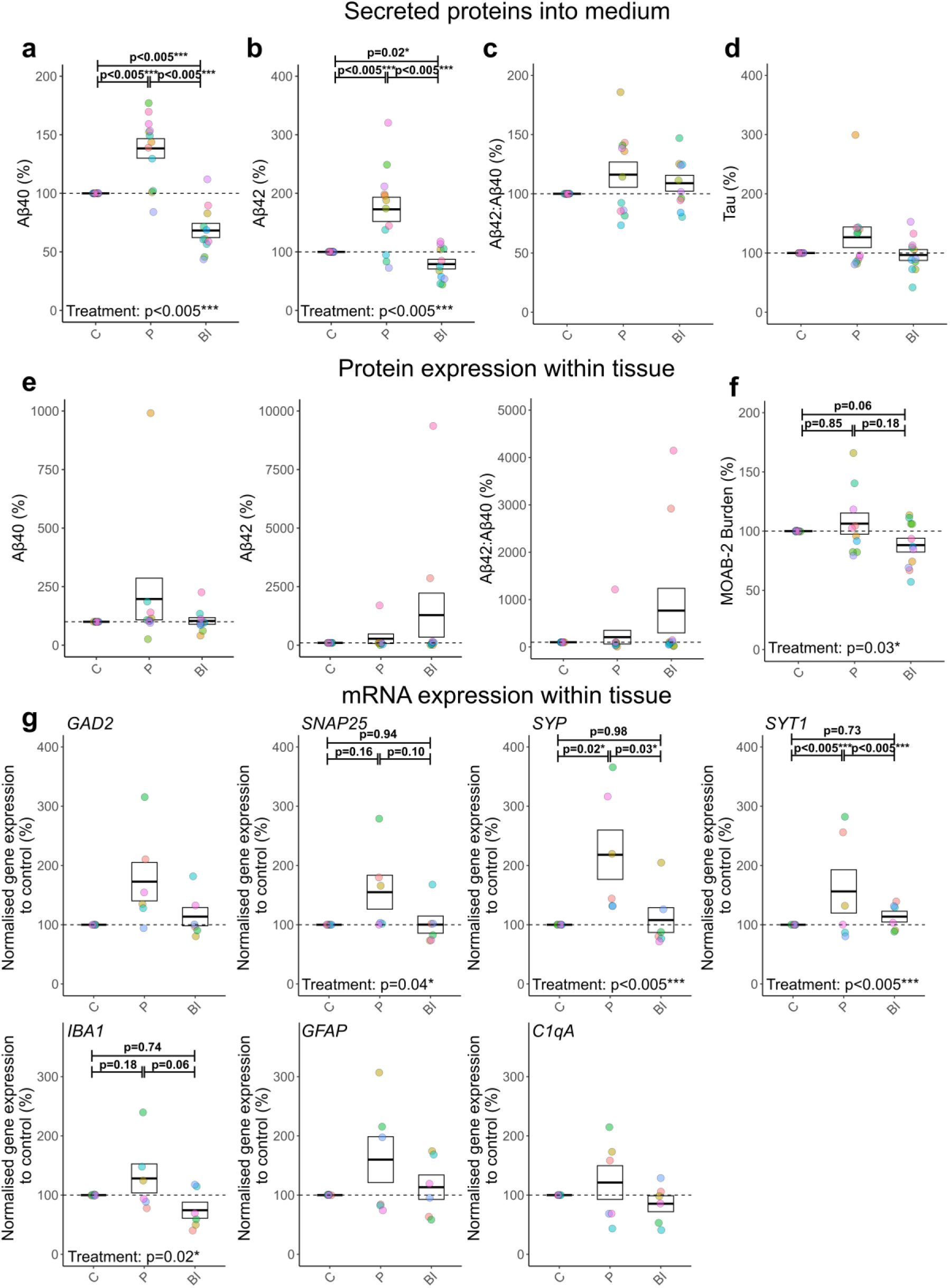
BACE1 inhibition and Phosphoramidon application for 7 div bi-directionally modulates Aβ production, altering synaptic gene expression. (a-g) C= Control, P= Phosphoramidon, BI= BACE Inhibitor. Box and dot plots showing normalised % change from control values when treated with Phosphoramidon or BACE1 inhibitor. Dots are coloured based on Case ID. Dashed line at 100% represents baseline (control), which samples were normalised to. Box represents standard error of the mean, with the thick line representing the mean. Statistics unless otherwise stated: GLMM with format var ∼ Drug treatment + Sex + (1|Age) + (1|Case ID). (a-d) Bidirectional modulation in response to Phosphoramidon and BACE1 inhibitor was seen with Aβ1-40 (gamma, χ²(2,36)=108.41, p<0.005***) (a) & Aβ1-42 (gamma, χ²(2,36)=53.37, p<0.005***) (b). There was no alterations on the levels of Aβ1-42/Aβ1-40 ratio (gaussian, χ²(2,36)=3.84, p=0.15) (c), and tau (log, χ²(2,36)=2.91, p=0.23) (d). Values taken from pg/mg concentration. (e) Protein expression within the slice, following guanidine extraction, assessed by ELISA. Aβ1-40 (Tukey, χ²(2,30)=3.20, p=0.20), Aβ1-42 (Tukey, χ²(2,30)=0.73, p=0.70), and Aβ1-42/Aβ1-40 (Tukey, χ²(2,30)=3.20, p=0.20) did not show significant differences in response to treatment with Phosphoramidon or BACE1 inhibitor. (f) MOAB expression showed significant alterations to treatment (gaussian, χ²(2,32)=6.73, p=0.03*), but did not show post hoc significant differences. (g) GAPDH normalised gene expression values (2∧ -ΔΔCT) in response to Phosphoramidon or BACE1 treatment from slices. Each graph shows gene expression normalised to the control treatment group. SNAP25 (Tukey, χ²(2,18)=6.37, p=0.04*), SYP (Tukey, χ²(2,18)=14.01, p<0.005***), SYT1 (gamma, χ²(2,18)=14.9, p<0.005***), and IBA1 (Tukey, χ²(2,18)=7.60, p=0.02*) all showed significant treatment effects. GAD2 (gaussian, χ²(2,18)=3.61, p=0.16), GFAP (Tukey, χ²(2,18)=1.75, p=0.42), and C1qA (gaussian, χ²(2,18)=2.72, p=0.26) did not show significant treatment effects.

### Exogenously applied AD-derived Aβ binds human post-synaptic structures

We next sought to examine the impact of disease-associated Aβ on synapses using sub-diffraction limit microscopy. Unlike many studies which use supraphysiological concentrations of synthetic Aβ, we sought to characterise the impact of pathophysiological levels of AD-derived Aβ (previously shown to contain highly bioactive low molecular weight Aβ oligomers^76^) on live adult human brain. HBSCs from the same patient were split into three conditions in a repeated measures design. Cultures were then exposed to either medium only control, medium containing 25% soluble AD-brain extract, (final concentration of 0.15 pM Aβ_1-42_, 7.05 pM Aβ_1-40_) (Aβ +ve) or medium containing 25% soluble AD-brain extract immunodepleted for Aβ (final concentration of Aβ_1-42_ undetectable by ELISA, <0.25 pM Aβ_1-40_) (Aβ -ve). HBSCs were exposed to treatments for 72 hours before collection for array tomography and Western blot analysis. Array tomography (**Fig. 4a**) revealed that HBSCs exposed to the Aβ +ve treatment showed a significant increase in the volume of oligomeric Aβ staining in the tissue when compared to Aβ -ve treatment (**Fig. 4b**). Interestingly, whilst there was no change in the level of oligomeric Aβ co-localising with the pre-synaptic marker synaptophysin (**Fig. 4c**), there was an increase in Aβ co-localising with the post-synaptic marker PSD95 (**Fig. 4d**), indicating a preferential binding to post-synaptic compartments in human cortical tissue. Interestingly, in the Aβ +ve group there was a decrease in synaptophysin density (**Fig. 4e**) but PSD95 density remained unchanged (**Fig. 4f**). Raw data, not normalised to medium-only control, is available in (**Supp. Fig. 3**). Neither Aβ +ve or Aβ –ve treatment resulted in loss of neuronal proteins Tuj1 or PGP 9.5 by Western blot (**Supp. Fig. 4**), indicating that soluble AD-brain extract did not result in wholesale loss of neurons. We also did not find any changes in the levels of synaptophysin, PSD95, or GFAP by Western blot between the Aβ +ve or Aβ –ve treatment groups (**Supp. Fig. 4**). However, there was a significant increase of PSD95 protein levels following soluble AD-brain extract treatment compared to medium control, perhaps indicating a compensatory response to the extract (**Supp. Fig. 4**). Overall, whilst the physical distribution of these proteins may change, the total level of proteins remains similar when comparing Aβ –ve and Aβ +ve conditions.

**Figure 4:**
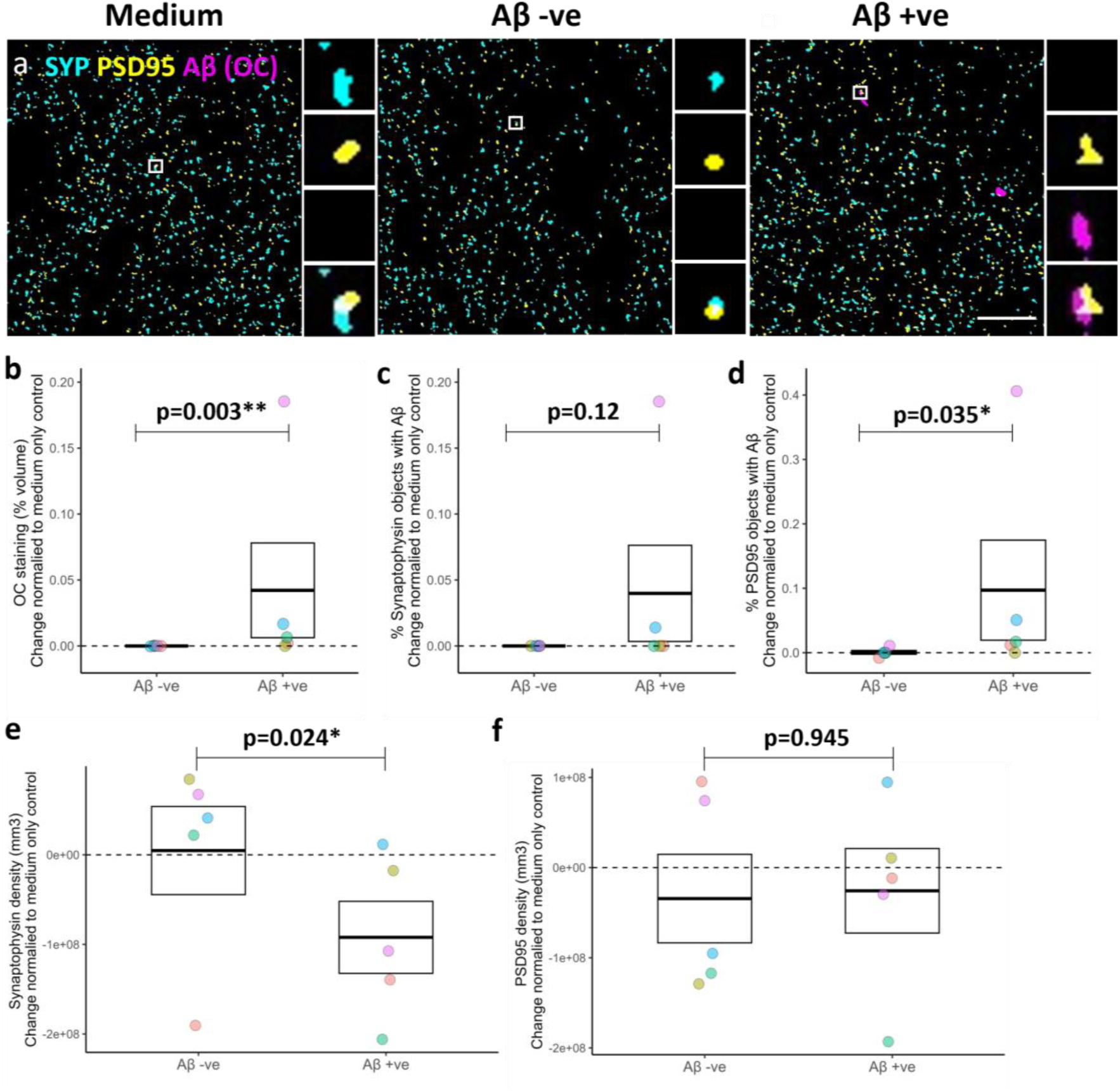
AD-derived Aβ colocalises to post-synapses and induces loss of synaptophysin in human brain slice cultures. **(a)** HBSCs were cultured for 72 hrs in either medium only, Aβ –ve soluble AD-brain extract or Aβ +ve soluble AD-brain extract. Slices were processed for array tomography and immunostained for pre-synapses (synaptophysin, SYP, cyan), post-synapses (PSD95, yellow), Aβ (OC, magenta). Single 70 nm segmented section shows representative staining and colocalization in each culture condition. Scale bar = 10 µm. Large boxes = 50 µm * 50 µm. Small boxes = 2 µm * 2 µm. **(b – d)** Slices cultured with Aβ +ve soluble AD-brain extract show an increase in the percentage volume of the 3D image stack occupied by OC (β = 0.56, 95% CI [0.186, 0.934], z.ratio = 2.936, p=0.003**) (B), no change in the percentage of pre-synapses containing Aβ (β = 0.182, 95% CI [-0.046, 0.41], z.ratio = 1.563, p = 0.118) **(c)** and an increase in the percentage of post-synapses with Aβ (β = 0.472, 95% CI [0.034, 0.911], z.ratio = 2.110, p = 0.0348*) **(d)**. Statistics: to ensure the assumptions of a statistical test were met, data was transformed to a binary readout. An increase in pathology was defined as >2 fold increase in staining, when compared to medium treated control, and analysed with GLMM using format var ∼ Treatment + (1|Case), family = binomial. **(e-f)** Culturing slices with Aβ +ve soluble AD-brain extract induced a loss of synaptophysin density (β = 501, 95% CI [89.8, 912], t(6.64) = 2.913, p = 0.024*) **(e)**. There was no effect on PSD95 density (β = 18.1, 95% CI [-573, 609], t(7.86) = 0.071, p = 0.945) **(f).** Statistics: LMEM performed on cube root transformed data var ∼ Treatment + (1|Case). To account for variability between cases, the difference between treatment groups and medium only controls were calculated. The dotted line at 0 represents change from medium only controls. Box represents standard error of the mean, with the thick line representing the mean. Data points refer to case means from image stacks. Dots are colored according to case ID.

### Aβ and tau pathological features are detected in live human brain samples from older patients, and are preserved in culture

Finally, we screened our HBSC samples for evidence of spontaneously occurring AD-relevant neuropathology. Whilst none of our donors had been diagnosed with a neurodegenerative disorder, 1 in 3 individuals are expected to develop dementia over their lifetime^77^, with pathological changes in the brain likely occurring decades before clinical disease presentation^78^. We therefore screened a subset of our cases (n=22) for AD-associated pathology. Immediately following tissue slicing, a subset of slices were fixed and immuno-stained for fibrillar-oligomeric Aβ (OC) and phosphorylated tau (phospho-tau Ser202/Thr205 (AT8)). Slices were then screened for pathological features and scored along the following criteria. For Aβ, we documented the presence or absence of extracellular Aβ plaques (**Fig. 5a**). For phospho-tau, we assessed whether we could observe neuropil thread-like staining or somatic tangle-like structures (**Fig. 5d**). Whilst we did detect Aβ staining in glial cells or blood vessels in a number of our samples, only 18% had extracellular OC staining typical of Aβ plaques (**Fig. 5b**). Phospho-tau staining revealed just under half of the cohort had neuropil thread-like staining, with 27% having evidence of neurofibrillary tangles (**Fig. 5e**). We found a significant effect of donor age on the presence of Aβ plaques (**Fig. 5c**) and on pathological phospho-tau (**Fig. 5f**), with greater levels of pathology seen in older donors. We did not detect Aβ plaques in any samples from donors under the age of 60, and no neurofibrillary tangles were detected in patients below the age of 50 years old. Having established that we could detect Aβ plaques in acute neocortical access tissue, we next sought to explore whether such features were observable, and retained over time in live HBSCs. We found that Aβ plaques could be labelled and live-imaged via multiphoton microscopy in HBSCs using methoxy-X04 (**Fig. 5g**) or Thioflavin-S (**Fig. 5h**) at 3 *div*. We were also able to observe Aβ plaques in proximity with MAP2 positive neurites and neuronal cell bodies in HBSCs fixed and immunostained after 7 *div* (**Fig. 5i,j**).

Following observations that pathological features were retained during the culture period, we then examined how the presence of Aβ or tau pathology in the slice impacted concentration of Aβ_1-40,_ Aβ_1-42_ or tau in the culture medium at 7 *div*. (**Fig. 6**). Whilst there was no relationship between plaque pathology and Aβ_1-40_ (**Fig. 6a**), Aβ_1-42_ (**Fig. 6b**) or total tau (**Fig. 6d**) concentration in the medium, there was a trend for increased Aβ_1-42_/Aβ_1-40_ ratio in samples with plaques (**Fig. 6c**). For tau pathology, there was similarly no effect of threads or tangles on Aβ_1-40_ (**Fig. 6g**), Aβ_1-42_ (**Fig. 6h**) or total tau (**Fig. 6j**) concentration in the medium, but there was a significant impact of pathology on the Aβ_1-42_/Aβ_1-40_ ratio (**Fig. 6i**). We did not find a relationship between the brain region taken, and the proportion of samples with either Aβ (**Fig. 6e**) or tau pathology (**Fig. 6k**). There was also no relationship observed between Aβ or tau pathology and patient *APOE* genotype, but we were limited by small group sizes for some *APOE* genotypes (**Fig 6 f, l**).

**Figure 5:**
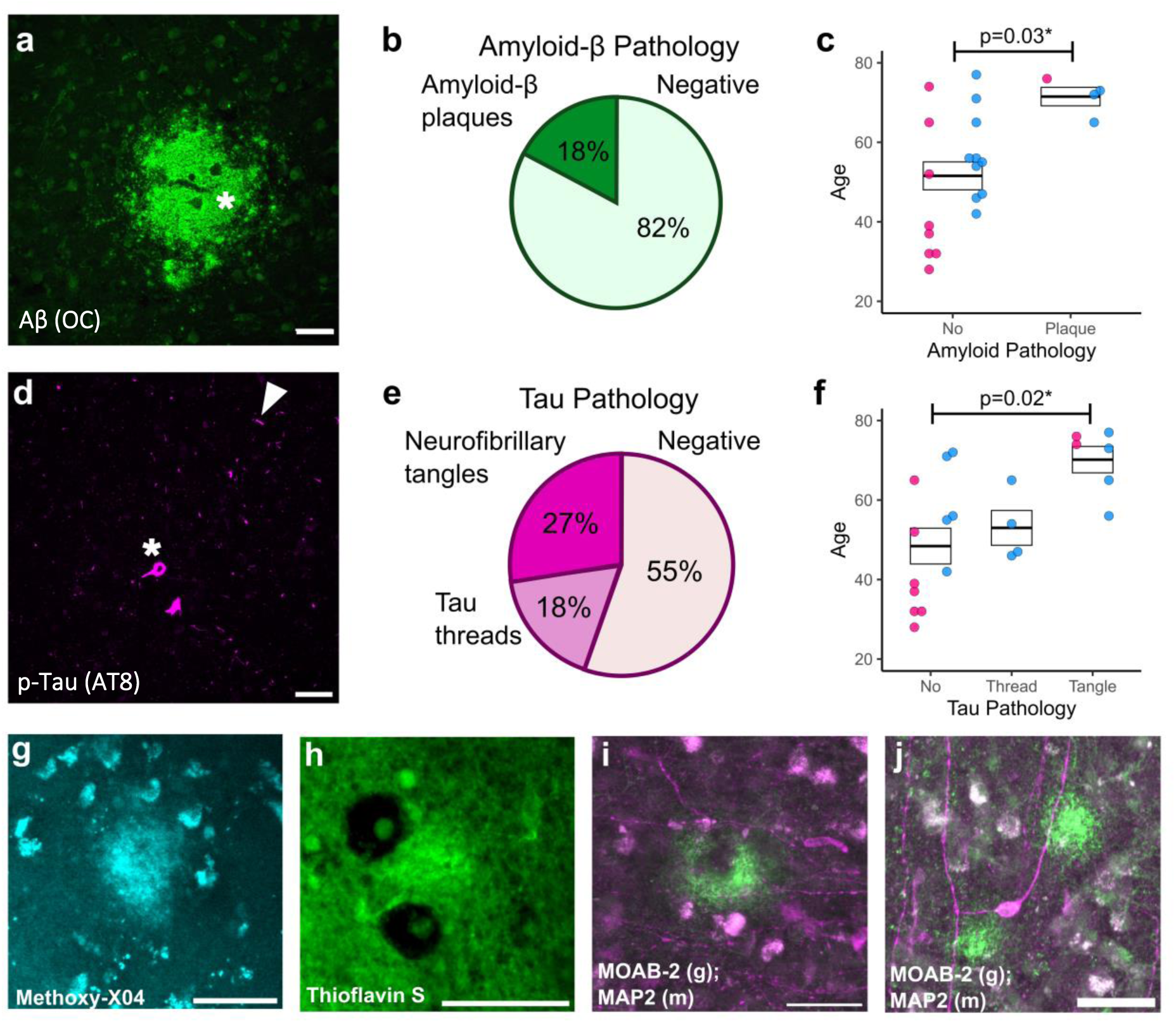
Endogenous AD-related pathology visible in human brain slices. **(a&d)** High magnification examples to highlight positive amyloid and tau pathology. **(a)** The asterisk denotes example Aβ plaque stained with OC. **(d)** The asterisk highlights neurofibrillary tangle pathology, and the arrowhead highlights thread pathology stained with AT8. **(b&e)** Pie charts highlighting the percentage of cases that have Aβ plaque pathology, determined by a visual inspection based on OC positive staining **(b)** or tau pathology, determined by a visual inspection based on AT8 staining **(e)**. **(c&f)** Box and dot plots showing the age of the patient compared to the Aβ (gaussian, F(1,22)=5.84, p=0.03*) **(c)** or tau (gaussian, F(2,22)=5.40, p=0.01**) **(f)** pathology score. A significant relationship between pathology and age is seen for Aβ and tau. Box represents standard error of the mean, with the thick line representing the mean. Statistics: LM with format Age ∼ Pathology score + Sex. **(g&h)** Live plaque imaging in culture using Methoxy-X04 or Thioflavin S. **(i&j)** Immunofluorescence staining following culturing using MOAB-2 for Aβ pathology (green) and MAP2 for neurons (magenta), showing Aβ plaques are retained in culture for at least 7 div. Scale bars are 50 µm. Blue dots represent males, pink dots represent females.

**Figure 6:**
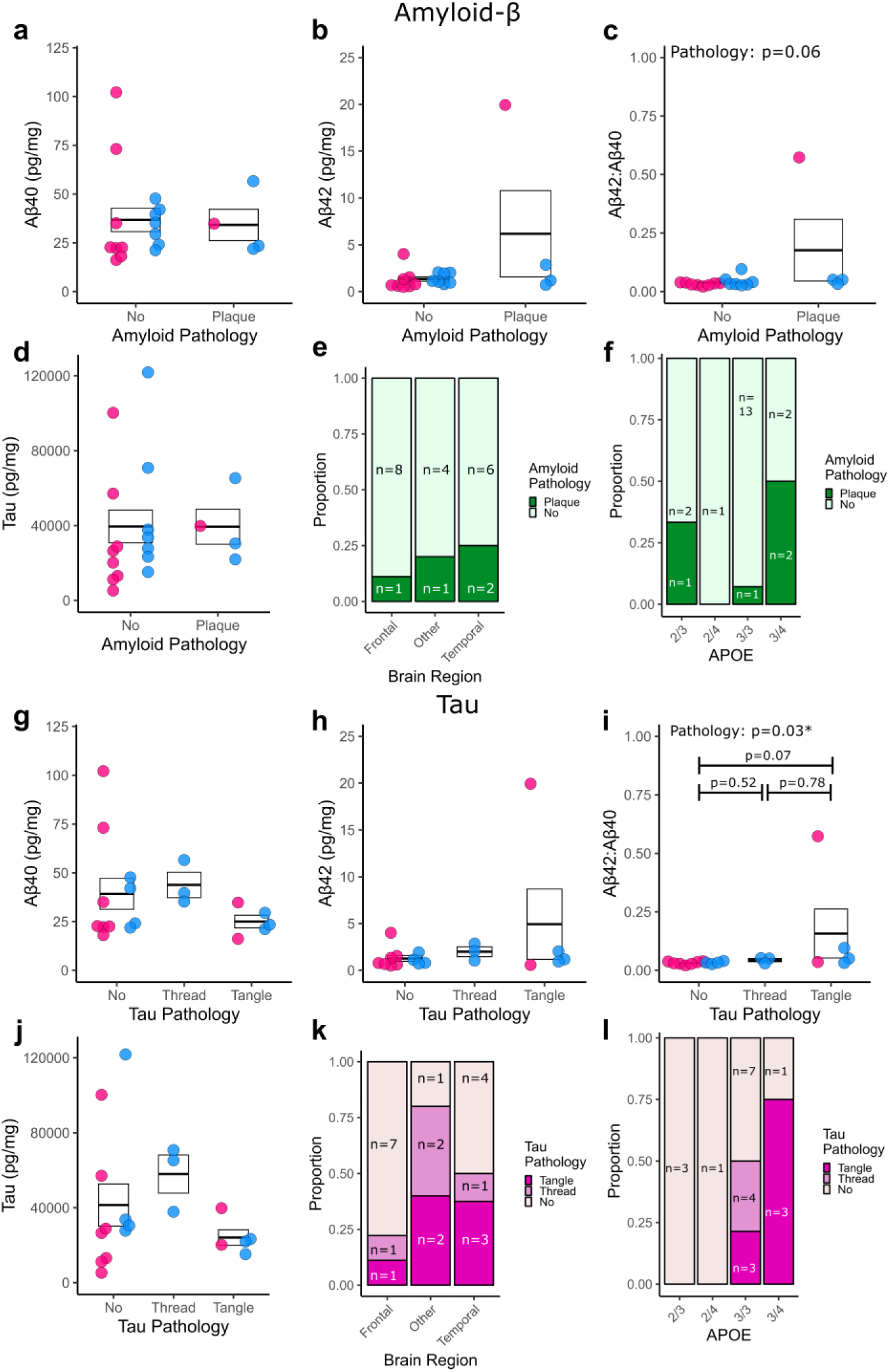
Assessment of relationship between Aβ or tau pathology in the slice compared with protein concentrations in the culture medium, brain region, and APOE genotype. Aβ (a-f) and tau (g-l) pathology scoring compared to Aβ1-40 (a&g), Aβ1-42 (b&h), Aβ1-42/Aβ1-40 (c&i), Tau (d&j), brain region (e&k), and APOE genotype (f&l). For Aβ pathology (a-f), there was no significant effect on Aβ1-40 (log, χ²(2,19)=1.57, p=0.21) (a), Aβ1-42 (Tukey, χ²(2,19)=1.67, p=0.20) (b) or tau (log, χ²(1,19)=0.07, p=0.80) (d) release into the medium. Aβ1-42/Aβ1-40 (Tukey, χ²(1,19)=3.64, p=0.06) (c) showed a trend towards plaque cases having higher ratios. There was no relationship between Aβ pathology and brain region (χ²(2,23)=0.56, p=0.75) (e), or APOE genotype (χ²(3,23)=4.55, p=0.21) (f). For tau pathology (g-l), there was no significant effect on Aβ1-40 (log, χ²(1,19)=4.84, p=0.09) (g), Aβ1-42 (Tukey, χ²(1,19)=3.84, p=0.15) (h) or tau (Tukey, χ²(1,19)=1.90,p=0.39) (j) release into the medium. There was a significant effect of tau pathology on Aβ1-42/Aβ1-40 ratio (Tukey, χ²(2,19)=7.27, p=0.02*) (i). There was no relationship between tau pathology and brain region (χ²(4,23)=5.18, p=0.27) (k), or APOE genotype (χ²(6,23)=9.10, p=0.17) (l). Statistics: GLMM with format var ∼ Pathology + Sex + (1|Age) or chi-squared test. Blue dots represent males, pink dots represent females.

## Discussion

To our knowledge, this study is the first to evaluate endogenous Aβ and tau dynamics, and spontaneous pathology in adult human brain slice cultures as well as to assess their response to pharmacological agents and human AD-derived Aβ. When examining endogenous protein concentrations in HBSC medium, there was no relationship between patient age and Aβ_1-42,_ Aβ_1-42_/Aβ_1-40_ ratio, or tau release into the medium. We did observe a trend towards reduced soluble Aβ_1-40_ with increasing patient age. Due to the lack of a blood-brain-barrier between the culture medium and the brain tissue, we believe HBSC medium is most comparable to *in vivo* cerebral ISF^79^. Whilst some studies have examined Aβ and tau turnover in human CSF using stable isotope labelling kinetics studies^24–28,80^, and direct sampling of ISF is regularly conducted in mice^81–84^, very few studies, to our knowledge, have directly sampled ISF levels of Aβ or tau in living human patients. One study linked increased Aβ levels with improved neurological status after acute brain injury^39^, whilst another found that ISF levels of Aβ_1-42_ were lowest in Normal Pressure Hydrocephalus (NPH) patients with cortical Aβ pathology compared to those without^40^. Whilst the brain injury study did not look at patient age as a variable, it may be that the higher levels of Aβ_1-40_ we see in HBSC medium from younger individuals reflects increased neuronal activity in these samples, either through baseline patient characteristics or tolerance to culture conditions. In contrast to the NPH study, we saw a trend towards increased Aβ_1-42_/Aβ_1-40_ in cases with plaque pathology compared to those without, with no effect on Aβ_1-40_ or Aβ_1-42_ levels. In addition, one case which had high Aβ plaque burden in the tissue, also had the highest levels of medium Aβ_1-42_ of the entire cohort (∼10 fold higher). Interestingly, we found a significant effect of tau pathology on the Aβ_1-42_/Aβ_1-40_ ratio in the culture medium, with samples containing tangles showing the highest Aβ_1-42_/Aβ_1-40_ ratio. More commonly conducted studies examining CSF from human patients have observed an overall decline in Aβ_1-42_ and Aβ_1-42_/Aβ_1-40_ ratio during normal ageing, which is exacerbated by both *APOE4* genotype and a diagnosis of AD^16–19,26,28,85^. Again, the lack of change in Aβ_1-42_ over time in our sample set differs from this, which may reflect different processes being captured in HBSC medium versus CSF/ISF, or could be due to limits in ELISA sensitivity for Aβ_1-42_. By contrast, our findings align well with studies in post-mortem brain tissue, that show the levels of soluble Aβ_1-40_ declines with age^36,37^. Further studies examining brain tissue, HBSC medium, ISF and CSF from the same patient donor could be highly informative in untangling how changes observed in HBSCs correlate with *in vivo* readouts.

Our finding that the levels of total tau in the HBSC medium were increased in temporal lobe samples when compared to frontal is of interest in the context of regional vulnerability to AD. Whilst four distinct patterns of tau spread have been reported in AD^66^, pathology generally originates in the medial temporal lobe^67,68^. Post-mortem brain studies report that, in healthy individuals, there is no difference in endogenous tau concentration between temporal and frontal lobes^86,87^. As we did not see an effect of age on tau concentration in HBSC medium, we hypothesise there could be regional differences in tau release in human brain tissue. Increased neuronal activity increases tau release in mice^88^, so it may be that there are baseline differences in neuronal activity or tau release in this region result that could contribute to greater vulnerability to tau pathology in later life. This observation, and future exploration of this hypothesis, is uniquely suited to HBSCs, as regional ISF measurements would be exceptionally challenging in human patients, and CSF measurements do not provide spatial information.

HBSCs facilitate testing potential therapeutics in live adult human brain tissue, as well as assessing the impact of increased disease-related protein levels-an impossible experiment in human patients. Human trials have shown that BACE1 inhibitors reduce plaque load, and lower CSF Aβ with little change to total or p-tau^58^. In line with this, we found that application of BACE1 inhibitor LY2886721 to HBSCs resulted in a significant decrease in Aβ_1-40_ and Aβ_1-42_ in HBSC medium with no change to tau levels. Previous studies have shown that administration of Phosphoramidon to mice or cell lines results in around a 2 fold increase in Aβ levels^71–74,89^. For the first time in live human tissue, we find that Phosphoramidon increases endogenous levels of Aβ and, results in increased levels of synaptic transcripts. Research suggests that elevated Aβ damages synapses^1^, but it’s crucial to highlight that most prior studies explore supraphysiological levels of exogenous Aβ, not physiological increases in endogenous Aβ. Indeed, evidence from mouse studies points to a hormetic effect of Aβ, where high levels of Aβ are synaptotoxic, but picomolar increases in Aβ often enhance synaptic function^3,90–96^. Our observation that increased endogenous Aβ concentration in HBSCs increases the expression of synaptic transcripts, provides evidence that a similar hormetic role of Aβ may be present in human brain tissue.

In contrast to manipulating endogenous production of Aβ in HBSCs, application of exogenous AD-brain derived Aβ resulted in loss of synaptic puncta. AD-brain derived Aβ oligomers, even at low picomolar concentrations as used here, have previously been shown to be toxic to synapses^1,41–44^. We provide the first direct evidence that Aβ oligomers preferentially bind to post-synaptic structures in live human brain tissue, in agreement with prior work in rat hippocampal neurons^97^ and rhesus macaque^45^. Whilst we find that oligomeric Aβ preferentially binds to post-synaptic structures, interestingly it is pre-synaptic puncta that are lost in our model. This builds on studies using free floating human brain slices that demonstrated synthetic Aβ accumulates in brain tissue and results in a loss of synaptophysin mRNA^98,99^. Other studies have previously shown that presynaptic proteins are lost first in response to amyloid pathology^98–101^, so it may be that despite Aβ being located post-synaptically, pre-synapses are more vulnerable to downstream pathological consequences. Studies in human post-mortem brain tissue have demonstrated that synaptic oligomeric Aβ and synapse loss is most prominent in the vicinity of Aβ plaques and accumulates to a greater degree in *APOE* ε4 carriers^51–53^, so future work exploring the impact of patient characteristics in response to Aβ challenge in HBSCs may help provide mechanistic insight into the relationship between *APOE* genotype and AD risk.

The fact that we can detect Aβ and tau pathology in a subset of our patient samples, and that these features are retained and can be live-imaged in culture, opens exciting future avenues to explore potential sporadic pre-clinical AD pathology in live adult human brain. Whilst there is some evidence that cetaceans, non-human primates, dogs and cats can develop spontaneous AD-like pathology^102–104^, wild-type mice do not develop pathology without the introduction of rare familial-AD mutations, often alongside overexpression of humanised APP or tau^105^. Compared with rodents, the human brain also contains seven times as many neurons and double the amount of synapses^11–14^. It seems clear then, especially for diseases such as Alzheimer’s disease, increased use of human model systems will improve our chances of translational success. Indeed, our previous work using HBSCs demonstrate that responses to pharmacological agents can differ between mouse and human brain slice cultures^62^. We also recently showed that oligomeric tau derived from brains of individuals with Progressive Supranuclear Palsy (PSP) can be taken up into synapses, and induced gliosis in HBSCs^63^. Other studies have established HBSCs as an excellent tool to investigate: physiological properties of human neurons^12,60,61,106–112^, tumour environments^113–115^, neurodegenerative disease^98,99,116–122^, epileptic activity^123,124^ and developmental disorders^125^. By correlating responses to treatments, as well as changes in endogenous dynamics, to patient age, sex, brain region, genetics and lifestyle risk factors, we propose that HBSCs represent an extremely powerful tool for both discovery and translational research.

## Extended Data

**Supplementary Figure 1:**
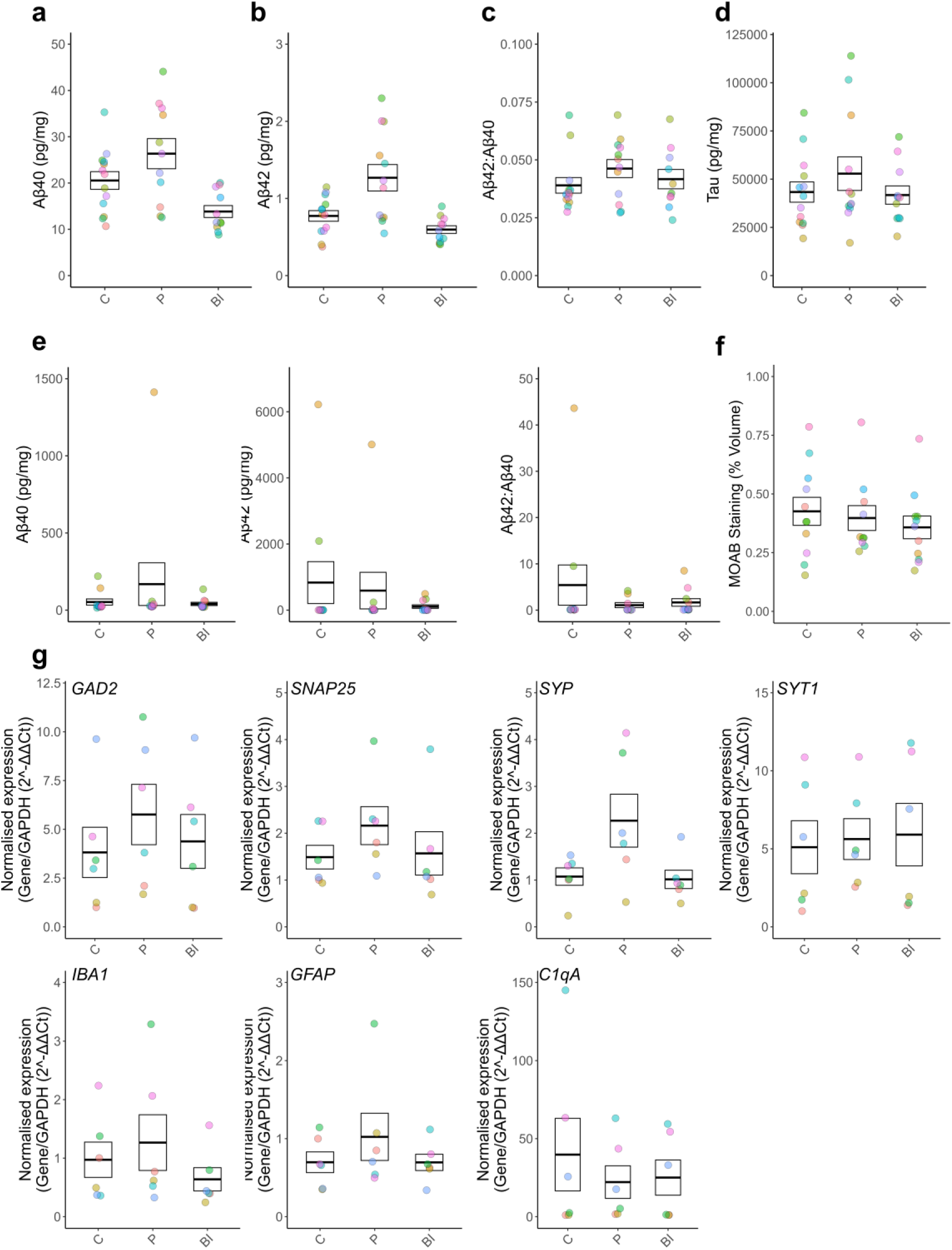
Raw values from Figure 3. **(a-g)** C= Control, P= Phosphoramidon, BI= BACE Inhibitor. Box and dot plots showing responses to treatment with Phosphoramidon or BACE1 inhibitor. Dots coloured based on Case ID. Box represents standard error of the mean, with the thick line representing the mean. **(a-d)** Responses to Phosphoramidon and BACE1 inhibitor treatment, as assessed by ELISA for Aβ1-40 **(a)**, Aβ1-42 **(b)**, Aβ1-42/Aβ1-40 ratio **(c)**, and tau **(d)**. **(e)** Protein expression for Aβ40, Aβ42, and Aβ1-42/Aβ1-40 within the slice, following guanidine extraction, assessed by ELISA. **(f)** MOAB staining, quantified as % volume of the imaged area. **(g)** GAPDH normalised gene expression values (GAD2, SNAP25, SYP, SYT1, IBA1, GFAP, C1qA) in response to Phosphoramidon or BACE1 treatment from slices.

**Supplementary Figure 2:**
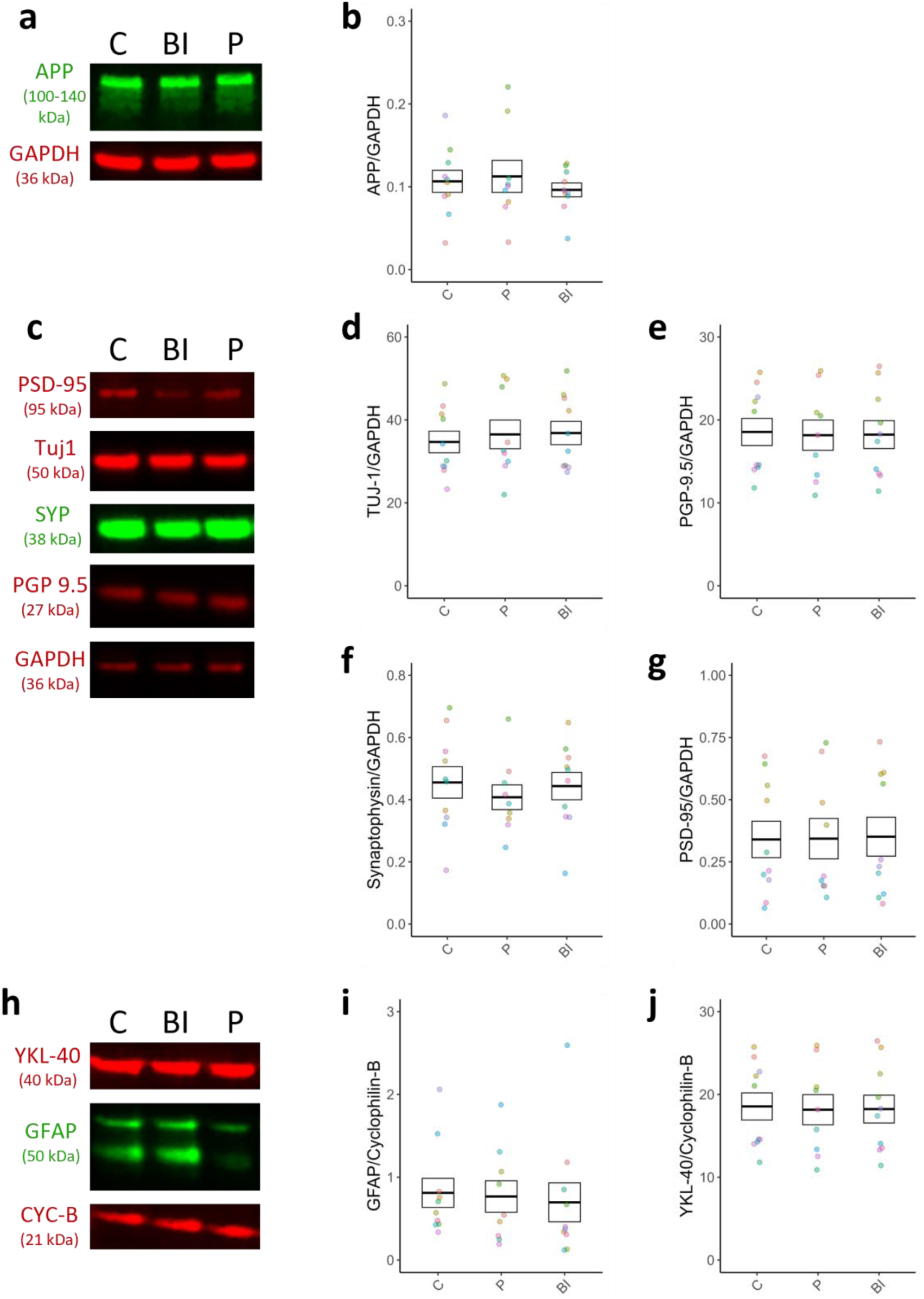
Protein expression changes following Phosphoramidon or BACE1 inhibitor treatment compared to medium control. C= Control, P= Phosphoramidon, BI= BACE Inhibitor. **(a)** Representative western blot and corresponding normalisation control (GAPDH) for APP, quantified in **(b)**. **(c)** Representative western blot and corresponding normalisation control (GAPDH) for APP, Tuj1, SYP, and PGP9.5. The quantification for this is shown in **(d-g)**. **(h)** Representative western blot images of YKL-40 and GFAP, and the corresponding normalisation control (CYC-B), quantified in **(i,j)**. Statistics: LM with format var ∼ Treatment + (1|Case). Box represents standard error of the mean, with the thick line representing the mean. Dots coloured based on Case ID.

**Supplementary Figure 3:**
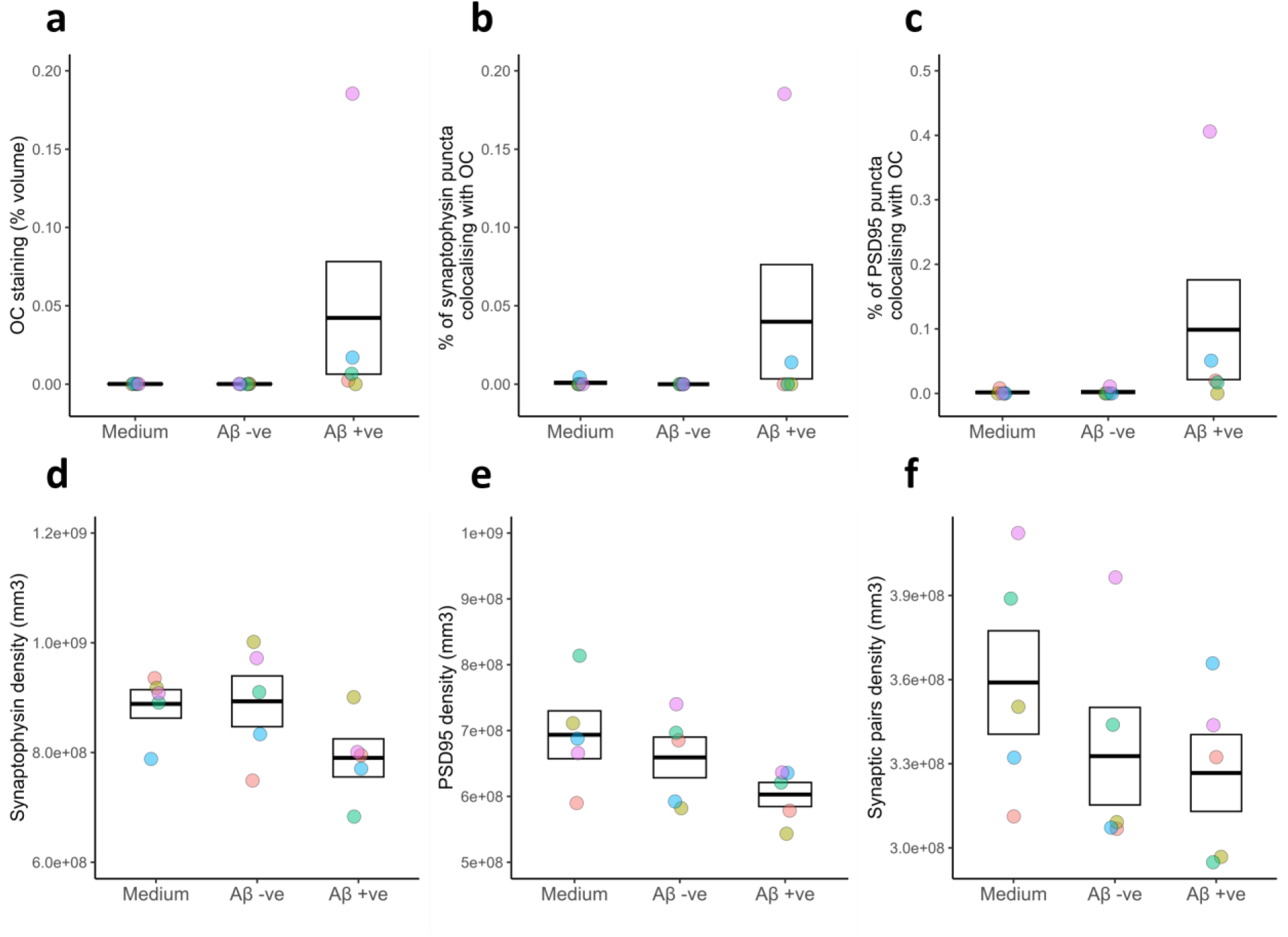
Raw values from array tomography data from. Figure 4**. (a-f)** Quantified responses to treatment with AD-brain extract compared to medium controls via array tomography **(a)** Percentage volume of the 3D image stack occupied by OC. **(b)** Percentage of pre-synapses containing Aβ. **(c)** Percentage of post-synapses containing Aβ. **(d)** Synaptophysin density, expressed as mm^3^. **(e)** PSD95 density, expressed as mm^3^. **(f)** Quantification of the density of synaptic pairs, as defined by a pre-synapse within 0.5 µm of a post-synapse. Box represents standard error and mean, with the thick line representing the mean., calculated from each image stack. Data points refer to case means. Dots coloured based on Case ID.

**Supplementary Figure 4:**
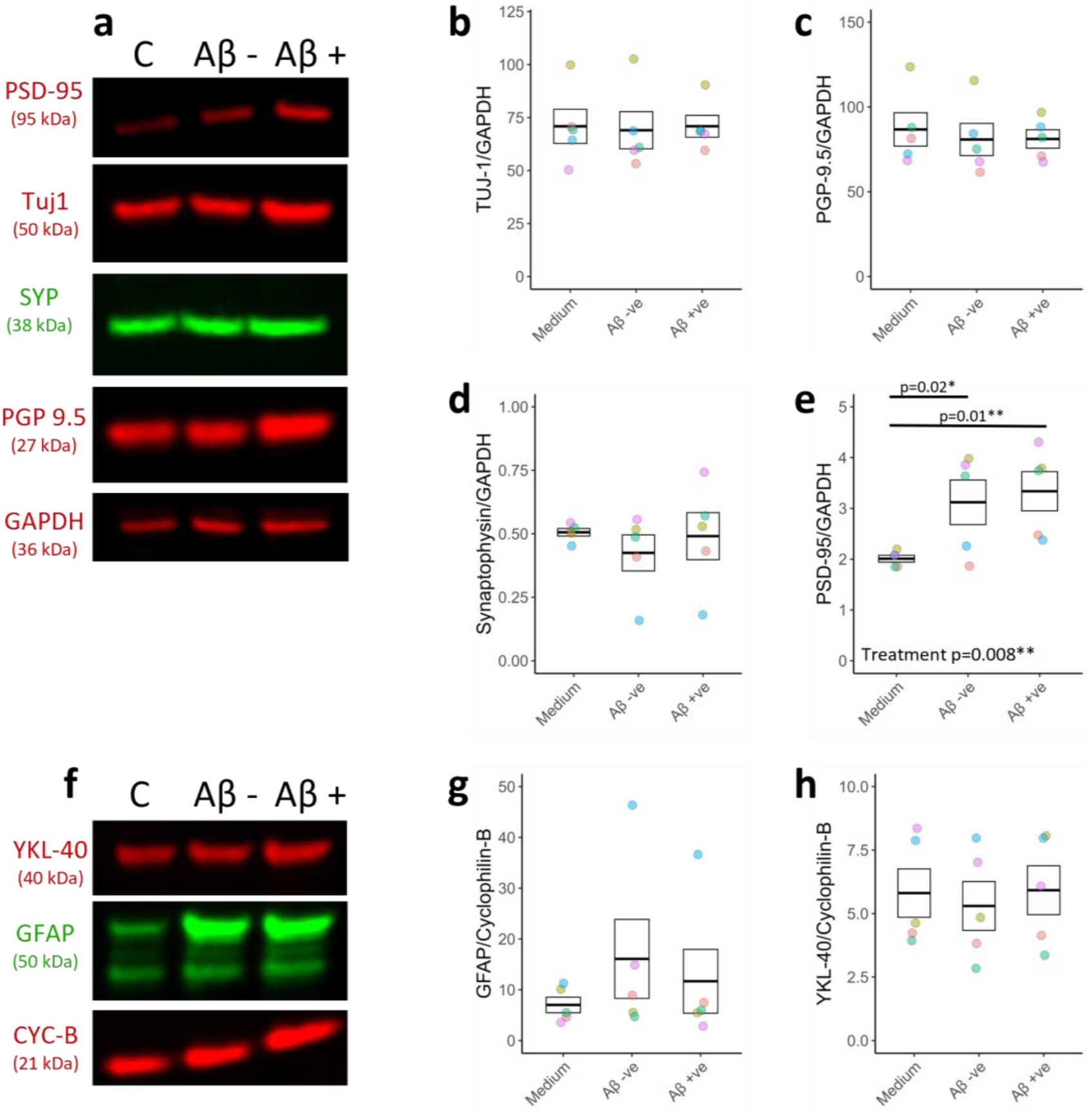
Protein expression changes following Aβ +ve or Aβ -ve AD-derived brain extract treatment compared to medium control. (a) Representative western blot and corresponding normalisation control (GAPDH) for PSD-95, Tuj1, SYP, PGP 9.5. The quantification for this is shown in **(b-e)**. **(e)** PSD-95 expression is significantly increased with AD-derived brain extract treatment compared to controls (linear, F(2,15)=9.25, p=0.008**). **(f)** Representative western blot images of YKL-40 and GFAP, and the corresponding normalisation control (CYC-B), quantified in **(g,h)**. C stands for control, Aβ - stands for Aβ immunodepleted AD-derived brain extract, Aβ + stands for mock immunodepleted AD-derived brain extract. Statistics: LM with format var ∼ Treatment + (1|Case). Box represents standard error of the mean, with the thick line representing the mean. Dots coloured based on Case ID.

## Methods

### Ethical approval and patient characteristics

Ethical approval for the use of surplus neocortical access tissue originating from patients undergoing tumour resection surgery was provided by the Lothian NRS Bioresource (REC number: 15/ES/0094, IRAS number: 165488, Bioresource No SR1319). NHS Lothian Caldicott Guardian Approval (Approval number CRD19080) was obtained to receive data on patient age (in years), sex, brain region provided and reason for surgery. Patients provided informed consent to donate surplus tissue (which would normally be discarded during surgery), and permit genetic testing of tissue, by signing the Lothian NRS Bioresource Consent form after discussion with a research nurse or neurosurgeon. Patient details are listed in **Table 1**. Post-mortem human temporal and frontal cortex tissue from people who died with Alzheimer’s disease was acquired from the Edinburgh Brain Bank and the Massachusetts Alzheimer’s Disease Research Center (grant 1P30AG062421-01). The Edinburgh Brain Bank is a Medical Research Council funded facility with research ethics committee approval (16/ES/0084). Experiments were approved by the Edinburgh Brain Bank ethics committee, the Academic and Clinical Central Office for Research and Development (ACCORD) and the medical research ethics committee AMREC a joint office of the University of Edinburgh and National Health Service Lothian, approval number 15-HV-016. The details of the post-mortem human cases are found in **Supplementary Table 1**. Inclusion criteria for cases included a clinical diagnosis of dementia and neuropathological diagnosis of Alzheimer’s disease with Braak stage V or VI. Exclusion criteria included substantial co-pathologies in the brain.

**Supplementary Table 1:**
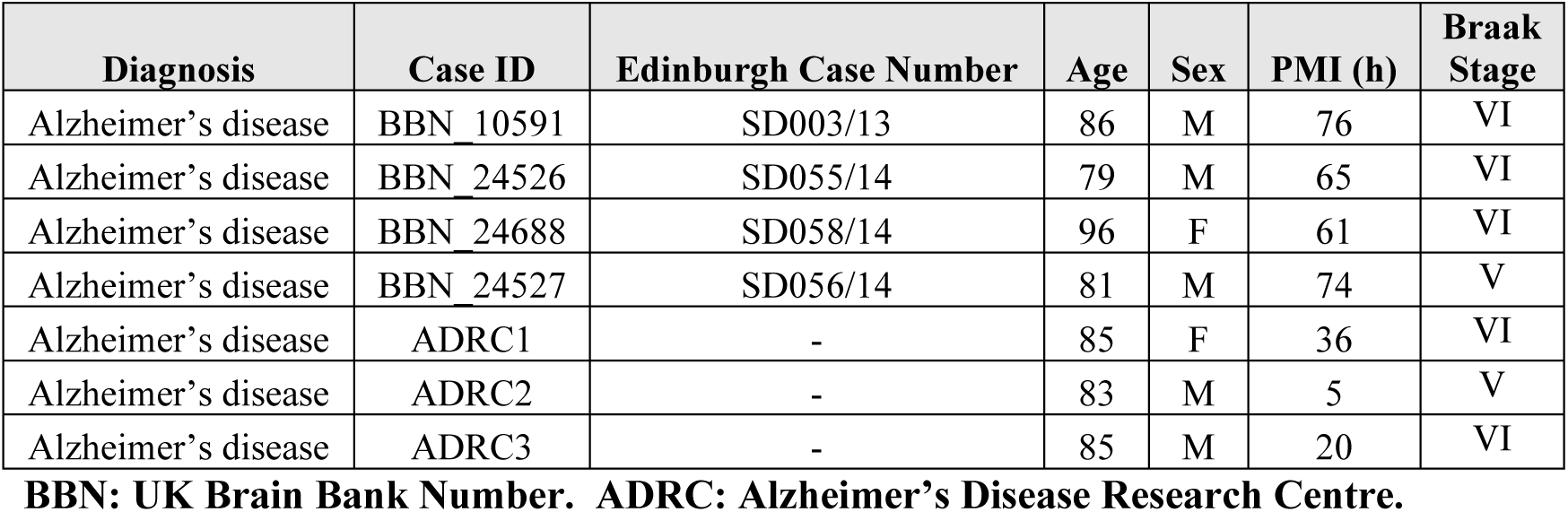
Human post-mortem case details.

### Generation and maintenance of human brain slice cultures (HBSCs)

Dissection and culture methods to generate human brain slice cultures (HBSCs) were adapted from published studies^60,106,126–128^. A summary schematic of tissue dissection is shown in **Fig. 1a**. Briefly, surplus, non-tumour, neocortical access tissue, was excised from patients undergoing brain tumour debulking surgery. This tissue was then immediately placed in ice-cold oxygenated artificial cerebrospinal fluid (aCSF) containing: 87 mM NaCl, 2.5 mM KCl, 10 mM HEPES, 1.62 mM NaH_2_PO_4_, 25 mM D-glucose, 129.3 mM sucrose, 1 mM Na-Pyruvate, 1 mM ascorbic acid, 7 mM MgCl_2_ and 0.5 mM CaCl_2_. aCSF is adjusted to a pH of 7.4 and ∼330 mOsm and sterilised by passing through a 0.22 μm filter before use. Thick sections of pia are removed from the excised tissue using ultra-fine forceps and the block is trimmed to permit optimal orientation to preserve cortical layering. Tissue is embedded in 2% agar, set briefly on ice, then mounted onto a Leica VT1200S vibratome specimen plate using Loctite superglue. Tissue was cut at <0.1mm/s speed in ice-cold oxygenated aCSF to generate 300 µm thick slices. Slices were further sub-dissected using a scalpel, to generate ∼3 mm wide columns containing all six cortical layers, and a small section of white matter. Once all sections were cut, slices were placed into a sterile wash solution (Wash Solution 1) composed of continuously-oxygenated Hanks Balanced Salt Solution (HBSS, ThermoFisher: 14025092), HEPES (20 mM) and 1X penicillin-streptomycin (ThermoFisher: 15140122), adjusted to pH 7.3, 305 mOsm, for 20 minutes at room temperature. Slices were then plated on 0.4 μm pore culture membranes (Millipore: PICM0RG50) placed inside 35 mm sterile culture dishes, containing 750 µl of Wash Solution 2. Wash Solution 2 was composed of 96% BrainPhys Neuronal Medium (StemCell Technologies: 5790), 1X N2 (ThermoFisher: 17502001), 1X B27 (ThermoFisher: 17504044), 40 ng/ml hBDNF (StemCell Technologies: 78005), 30 ng/ml hGDNF (StemCell Technologies: 78058), 30 ng/ml Wnt7a (Abcam: ab116171), 2 μM ascorbic acid, 1 mM dibutyryl cAMP (APExBIO: B9001), 1 ug/ml laminin (APExBIO: A1023), 1X penicillin/streptomycin (ThermoFisher: 15140122), 3 units/ ml nystatin (Merck: N1638) and 20 mM HEPES. For all samples, the time taken from surgical tissue removal to slices being plated in Wash Solution 2 was kept to a minimum (< 2 hours). Slice cultures were kept in Wash Solution 2 in an incubator at 37°C with 5% CO_2_ for between 1-5 hours. Following this, Wash Solution 2 was aspirated and replaced with pre-warmed sterile Maintenance Medium (identical composition to Wash Solution 2, but with HEPES removed). After plating, 100% Maintenance Medium replacement occurred twice weekly at 4 DIV (days *in vitro*) and 7 DIV (unless otherwise specified). Culture Medium was collected and flash frozen on dry ice at 7 DIV for further analysis.

### HBSC drug treatments

For the application of Aβ-modulating pharmaceuticals, 5 mM BACE1 inhibitor LY2886721 (Abcam: ab223886) and 100 mM Phosphoramidon (Sigma-Aldrich: R7385) stock solutions were generated by dissolving drugs in DMSO (Sigma-Aldrich: D2438) and sterile Milli-Q water respectively. Three treatment groups were allocated per human case, with a control condition having no treatment (i.e. 100% Maintenance Medium only). The respective drug treatments were added at 1/1000 dilution into Maintenance Medium to produce a final concentration of 5 µM BACE inhibitor or 100 µM of Phosphoramidon. 2 μl of medium (either control, BACE-1 inhibitor or Phosphoramidon treated) was added on top of each slice to ensure complete penetration of treatments through the slice tissue. Treatment was applied for 7 DIV, following which medium and slices were collected and processed as required.

### AD-brain soluble extract preparation

Soluble extracts from AD frontal and temporal cortex from Alzheimer’s patients detailed in **Supplementary Table 1** was generated as described previously^129^. Frozen samples were thawed on ice, finely diced and homogenised in aCSF (124 mM NaCL, 2.8 mM KCl, 1.25 mM NaH_2_PO_4_, 26 mM NaHCO_3_), pH 7.4, supplemented with Complete Mini EDTA-free protease inhibitor cocktail tablets (Roche, 11836170001), using a Dounce homogenizer. Homogenised tissue was then transferred to 15 mL protein LoBind® tubes (Eppendorf, 0030122216), placed on a roller for 30 minutes at 4°C, and then centrifuged at 2000 x g at 4°C for 10 minutes to remove insoluble debris. The supernatant was removed and ultracentrifuged at 4°C for 110 minutes at 200,000 x g. The resulting supernatant, the soluble protein extract (s-extract), was dialysed in aCSF (as above) at 4°C for 72 hours (Slide-A-Lyzer G2 Dialysis Cassettes) to remove salts, impurities and any pharmacological agents the donor may have been exposed to. The aCSF was replaced every 24 hours. Following dialysis, the s-extract was pooled and divided in two for immunodepletion of total Aβ. The two portions of s-extract were incubated overnight at 4°C with Protein A agarose (PrA) beads (30μL / ml of sample) (ThermoFisher Scientific, 20334) with either anti-β-amyloid 17-24 (4G8, mouse IgG2b, 1:100, BioLegend, #SIG-39200) and anti-β-amyloid 1-16 (6E10, mouse IgG1 1:100 BioLegend, #SIG-39320) to generate “Aβ negative” s-extract, or mock-immunodepleted with an anti-GFP antibody (mouse IgG, 1:50, DSHB, DSHB-GFP-8H11) to create the “Aβ positive” s-extract. The s-extracts were then centrifuged at 2500 x g for 5 minutes, the supernatant was collected into fresh protein LoBind® tubes and the process repeated two times (to result in a total of three overnight immunodepletion steps, each time the s-extract being exposed to fresh antibody and bead solutions). Finally, the Aβ negative and Aβ positive s-extract supernatants were collected into fresh protein LoBind® Eppendorf’s and stored at -80 °C until needed. The concentration of Aβ_1-40_ and Aβ_1-42_ in the s-extracts was determined by commercially available ELISA kits (ThermoFisher Scientific: KHB3481 and KHB3544). Aβ_1-42_ was undetectable in the Aβ negative sample, whilst it was 0.6 pM in the Aβ positive s extract. Aβ_1-40_ was <1 pM in the Aβ negative s-extract, whilst it was 28.2 pM in the Aβ positive extract.

### HBSC AD-brain soluble extract treatment

HBSCs from the same patient were split into three treatment groups in a repeated measures design. Group 1 was treated with 100% medium (control), Group 2 was treated with 75% medium: 25% Aβ negative s-extract (described above) (Aβ negative), Group 3 was treated with 75% medium: 25% Aβ positive s-extract (described above) (Aβ positive). 2 μl of treated culture medium was added on top of each slice to ensure complete exposure to treatments. Slices were collected after 3 DIV and either collected for western blot or fixed for array tomography processing (below).

### Array Tomography

Slices were processed for array tomography as previously described^130^. Briefly, following collection, slices were fixed in 4% paraformaldehyde for 1.5 hours, dehydrated in ethanol and incubated in LR White resin (Agar Scientific: AGR1281) overnight at 4°C. Individual samples were cured in LR White in gelatin capsules overnight at 53°C. Samples were cut into ribbons of ultrathin serial 70 nm sections using a histo jumbo diamond knife (DiATOME, agar scientific) and an Ultracut (Leica EM UC7). Ribbons were mounted on glass coverslips (treated with fish-skin gelatin), dried on a slide warmer and outlined with hydrophobic pen. Ribbons were rehydrated in 50 mM glycine in TBS and blocked in a solution of 0.1% fish skin gelatin and 0.05% Tween in TBS for 45 minutes at room temperature. Ribbons were incubated with primary antibodies (see **Supplementary Table 2**) at room temperature for 2 hours. Ribbons were washed with TBS, and secondary antibodies (see **Supplementary Table 2**) were applied for 45 minutes at room temperature. Ribbons were then washed with TBS. Finally, ribbons were mounted on glass slides with ImmuMount and imaged within 72 hours on a Zeiss Axio imager Z2.

**Supplementary Table 2:** Antibody details for array tomography.

| Antibody | species / isotype | product code | concentration | secondary | concentration | product code |
| --- | --- | --- | --- | --- | --- | --- |
| PSD95 | GP | synaptic systems: 124-014 | 1:50 | Donkey anti-guinea pig 488 | 1:50 | 706-545-148 |
| OC (fibrillar-oligomeric Aβ) | Rabbit Serum | Merck: ab2286 | 1:200 | Donkey anti-rabbit IgG 594 | 1:50 | A21447 |
| Synaptophysin | Goat IgG | AF555 | 1:50 | Donkey anti-goat 647 | 1:50 | A21447 |

### Microscopy and image analysis for array tomography

For array tomography imaging (**Fig. 4**), experimenters were blinded to case information during image processing and analysis. Images were acquired using Zen software using a 63x oil immersion objective (1.4 numerical aperture) on a Zeiss AxioImager Z2 with a CoolSnap digital camera. Two regions of interest (ROIs) were chosen per slice and imaged in the same location on 15-30 sequential sections. Imaging parameters were kept the same, and an image was taken of the negative control in each session to ensure there was no non-specific staining. Individual images from each ROI were combined into a 3D image z-stack and a median background filter was applied in Image J with custom batch macros. Using custom MATLAB scripts, image stacks were aligned using rigid registration. Each channel was segmented using an auto-local thresholding algorithm to binarize images and remove any objects present in only a single section (noise). Segmented images were run through custom MATLAB and Python scripts to determine object density, colocalization between channels, and the burden of the staining (% volume of 3D image stack occupied by the stain). Objects were considered colocalized if at least 25% of the 3D volume overlapped. For pre-and post-synaptic objects to be considered a synaptic pair, the distance between the centre of each object had to be ≤0.5 μm. Saturation was minimized during image acquisition and only applied for figure visualization. All custom software scripts are available on GitHub https://github.com/Spires-Jones-Lab. For more details, please see our methods video demonstrating this technique at https://doi.org/10.7488/ds/297.

### Immunofluorescence

For standard immunofluorescence techniques (**Fig. 1**, **Fig. 5**). Following culturing, slices were transferred to 4% PFA in PBS overnight and then stored in PBS until needed. Slices were rinsed in PBS with 1% Triton-X for 10 minutes. For Iba1, NeuN and MAP2, slices underwent an additional antigen retrieval step (10 minutes in a pressure cooker in IHC Antigen Retrieval Solution (Invitrogen: 00-4956-58), followed by cooling in ice for 30 minutes) before commencing the following protocol. Slices were transferred into 70% ethanol for 5 minutes. Slices were placed into autofluorescence eliminator reagent (Chemicon: 2160) for 5 minutes. Slices were then transferred into 70% ethanol for 5 minutes. All the following steps were performed on a shaker. Slices were rinsed 3 times for 10 minutes in PBS with 1% Triton-X, followed by a 1-hour block in normal goat/donkey serum (1:200, diluted in PBS with 1% Triton-x). Slices were then incubated in primary antibody (details listed in **Supplementary Table 3**) overnight at 4 °C. Following this, slices were rinsed in PBS with 1% Triton-x 3 times. Slices were then incubated in the corresponding secondary antibody (details listed in **Supplementary Table 3**) at a concentration of 1:1000 in PBS with 1% Triton-x overnight at 4 °C. Slices were finally rinsed 3 times for 10 minutes in PBS. Slices were then mounted with fluoroprotectant mountant medium (Vectashield: H-1000) and imaged.

**Supplementary Table 3:** Antibody details for immunofluorescence.

| Antibody | species<br>/<br>isotype | product<br>code | concentration | secondary | concentration | product<br>code |
| --- | --- | --- | --- | --- | --- | --- |
| AT8 (p-tau Ser202/Thr205) | mouse IgG1 | MN1020 | 1:1000 | Donkey anti-mouse AF 405 | 1:1000 | AB175658 |
| OC (fibrillar-oligomeric Aβ) | Rabbit Serum | Merck: ab2286 | 1:1000 | Donkey anti-rabbit AF 594 | 1:1000 | A21447 |
| MOAB-2 (pan-Aβ) | Mouse | Millipore: MABN254 | 1:1000 | Goat anti-mouse AF594 | 1:1000 | A21203 |
| MAP2 | Guinea pig | Synaptic Systems: 188004 | 1:1000 | Goat anti-guinea pig AF 488 | 1:1000 | A11073 |
| NeuN | Rabbit | Invitrogen: 702022 | 1:500 | Donkey anti-rabbit AF 488 | 1:1000 | A21206 |
| Iba1 | Rabbit | WAKO: 019-19741 | 1:1000 | Donkey anti-rabbit AF 488 | 1:1000 | A21206 |
| GFAP | Chicken | Abcam: AB4674 | 1:1000 | Goat anti-chicken AF405 | 1:1000 | A48260 |
| P2YR12 | Rabbit | Atlas antibodies: HPA014518 | 1:1000 | Goat anti-rabbit AF 647 | 1:1000 | A21244 |

### Microscopy and image analysis for immunofluorescence

Slices were imaged on a multiphoton microscope (Leica TCS SP8; Coherent Chameleon laser at 760 nm), using LAS X software, under a 25x objective. For pathology assessment (**Fig. 5**) whole slices were imaged and assessed for Aβ or tau pathology blinded to experimental condition of patient details. Immediately following tissue slicing, a subset of slices were fixed and immuno-stained for fibrillar-oligomeric Aβ (OC) and phosphorylated tau (phospho-tau Ser202/Thr205 (AT8)) (**Fig. 5a**). Slices were then screened for pathological features and scored along the following criteria. For Aβ, we documented the presence or absence of extracellular Aβ plaques (**Fig. 5b**). For phospho-tau, we assessed whether we could observe neuropil thread-like staining or somatic tangle-like structures (characteristic flame shaped AT8 positive inclusions) (**Fig. 5c**). For assessing total Aβ burdens within the slice (using MOAB-2, a pan-Aβ that does not detect APP) (**Fig 3**), a 50 µm stack from cortical layers I/II, III/IV, V/VI and white matter was taken in regions free from Aβ plaques. Each channel was segmented using an auto-local thresholding algorithm to binarize images. Segmented images were then run through custom MATLAB script to determine object density the burden of the staining (% volume of 3D image stack occupied by the stain). All custom software scripts are available on GitHub https://github.com/Spires-Jones-Lab. The MOAB-2 burden across all regions was averaged per slice to produce the final “slice burden”.

### Plaque imaging in live HBSCs

HBSCs were imaged on a multiphoton microscope (Leica TCS SP8; Coherent Chameleon laser at 760 nm), using LAS X software, under a 25x objective in a heat chamber (Oko Lab) set to 37 °C. Slices were imaged in Maintenance Medium to prevent changes in osmolarity or pH affecting live imaging readouts. Slices were pre-treated on top with either 1% Thioflavin-S diluted in Maintenance Medium or 100 µM Methoxy-X04 diluted in PBS for at least 20 minutes before imaging. Slices were imaged for a maximum of one hour, following which they were returned to the incubator for further culturing.

### Electrophysiology recordings

Whole-cell patch clamp recordings from 7 DIV HBSCs were made on a Scientifica based recording rig, with recordings amplified with a MultiClamp 700B amplifier (Axon Instruments), and digitised using a Digidata 1550B using pClamp 11.2. Slices were removed from culture membrane and placed into aCSF (33°C) made of: 125 mM NaCl, 2.5 mM KCl, 25 mM NaHCO_3_, 1.25 mM NaH_2_PO_4_, 25 mM Glucose, 1 mM MgCl_2_, 2 mM CaCl_2_, 1 mM Na-Pyruvate, 1 mM Na-Ascorbate. Following this, cells were targeted for whole-cell patch clamp recordings. Electrodes of 4-6 MΩ resistance were filled with a Potassium Gluconate based internal solution, made of in mM: 142 K-Gluconate, 4 KCl, 2 MgCl_2_, 0.1 EGTA, 10 HEPES, 2 Na_2_-ATP, 0.3 Na_2_-GTP, 10 Na_2_-Phosphocreatine. Biocytin was also included at a concentration of 0.1%, and the solution was pH 7.35, between 290 and 310 mOsm. Cells were recorded in gap free in current clamp for spontaneous synaptic currents and hyperpolarising and depolarising steps were elicited to evoke responses from the neuron.

### APOE Genotyping

Samples were genotyped by the Edinburgh Genetics Core. All samples were genotyped using the TaqMan SNP Genotyping Assays. Taqman genotyping was carried out on the QuantStudio12KFlex to establish APOE variants using the following assays: C__3084793_10 for rs429358 and C__904973_10 for rs7412.

### ELISA

HBSC medium was collected after 7 DIV, flash frozen on dry ice in LoBind® Eppendorf’s and frozen until use. Total protein was measured using a micro BCA assay according to manufacturers instructions (ThermoFisher Scientific: 23235). ELISAs for Aβ_1-40_ (ThermoFisher Scientific: KHB3481), Aβ_1-42_ (ThermoFisher Scientific: KHB3544) and total tau (ThermoFisher Scientific: KHB0042) were run according to manufacturers instructions. Culture medium was run undiluted for Aβ_1-40_ and Aβ_1-42_, whilst medium was diluted 1/500 in standard dilution buffer (provided with ELISA kit) before tau analysis. Values were obtained by comparing to a standard curve (pg/ml) then sample values were normalised to the total medium protein content (from BCA assay) to provide pg/mg. To assess the levels of Aβ in the slice tissue, slices were homogenised in 5M Guanidine Hydrochloride (Sigma #G3272 ;20 μl/ slice) for 4 hours at room temperature. 180 μl / slice ice cold PBS supplemented with 1x Halt Protease Inhibitor Cocktail (ThermoFisher Scientific: 78429) was then added to the guanidine extract, then spun for 20 minutes at 4°C at 16,000 x g. The supernatant was then run diluted (1/1 to 1/100) on the ELISA to accommodate differences in sample concentrations of Aβ. Values were obtained by comparing to a standard curve (pg/ml) then sample values were normalised to the total slice protein content (from BCA assay) to provide pg/mg.

### RNA isolation and quantitative reverse transcription PCR (RT-qPCR)

HBSCs (8 pooled slices) were removed from the culture membrane AT 7 DIV with a scalpel blade into pre-filled bead mill tubes (Fisherbrand: 15555799) containing RNA*later* stabilisation solution (ThermoFisher: AM7020) and kept at 4°C until RNA extraction. RNA was extracted using the TRIzol Plus PureLink RNA Purification Kit (Invitrogen: 12183555) which is a mixture of guanidinium thiocyanate-phenol-chloroform extraction and silica-cartridge purification methods. After removal of RNA*later*, samples were homogenised in 1 mL of TRIzol using a Bead Mill 24 Homogeniser (Fisherbrand: 15515799) to ensure equal homogenisation. Since HBSCs have high fat content, the lysates were centrifuged for 5 mins at 13,000 rpm at 4°C to pellet fatty tissue and cell debris, then the clear supernatant was transferred to a new tube for processing. RNA was quantified using a NanoDrop 2000 spectrophotometer (ThermoFisher: ND2000). To remove DNA contamination before qPCR, RNA samples were treated with Turbo DNAse (Invitrogen: AM2238) and the reaction was stopped with UltraPure EDTA (ThermoFisher: 15575020) at a final concentration of 15 mM. Quantitative RT-PCR was performed in 20 μl reactions in 96-well plates (ThermoFisher:AB0800W) using BRYT Green Dye contained in the GoTaq 1-Step RT-qPCR kit (Promega: A6020) in the CFX96 Touch Real-Time PCR Detection system (Bio-Rad). We used 100 ng of RNA per reaction and added extra MgCl_2_ (ThermoFisher: R0971) at a final concentration of 3.25 mM (including the 2 mM MgCl_2_ contained in the Master Mix), to each reaction to counteract EDTA chelation from the DNAse treatment (EDTA concentration in each qPCR reaction was 3.75 mM). Primers (ThermoFisher) (detailed in **Supplementary Table 4**) were used at a final concentration of 250 nM. Ct values were normalized to the housekeeping gene GAPDH, which was amplified in parallel. Melt curves were checked to ensure correct amplification of a single PCR product. The 2∧-ΔΔCT method was utilized to calculate relative gene expression levels.

**Supplementary Table 4:**
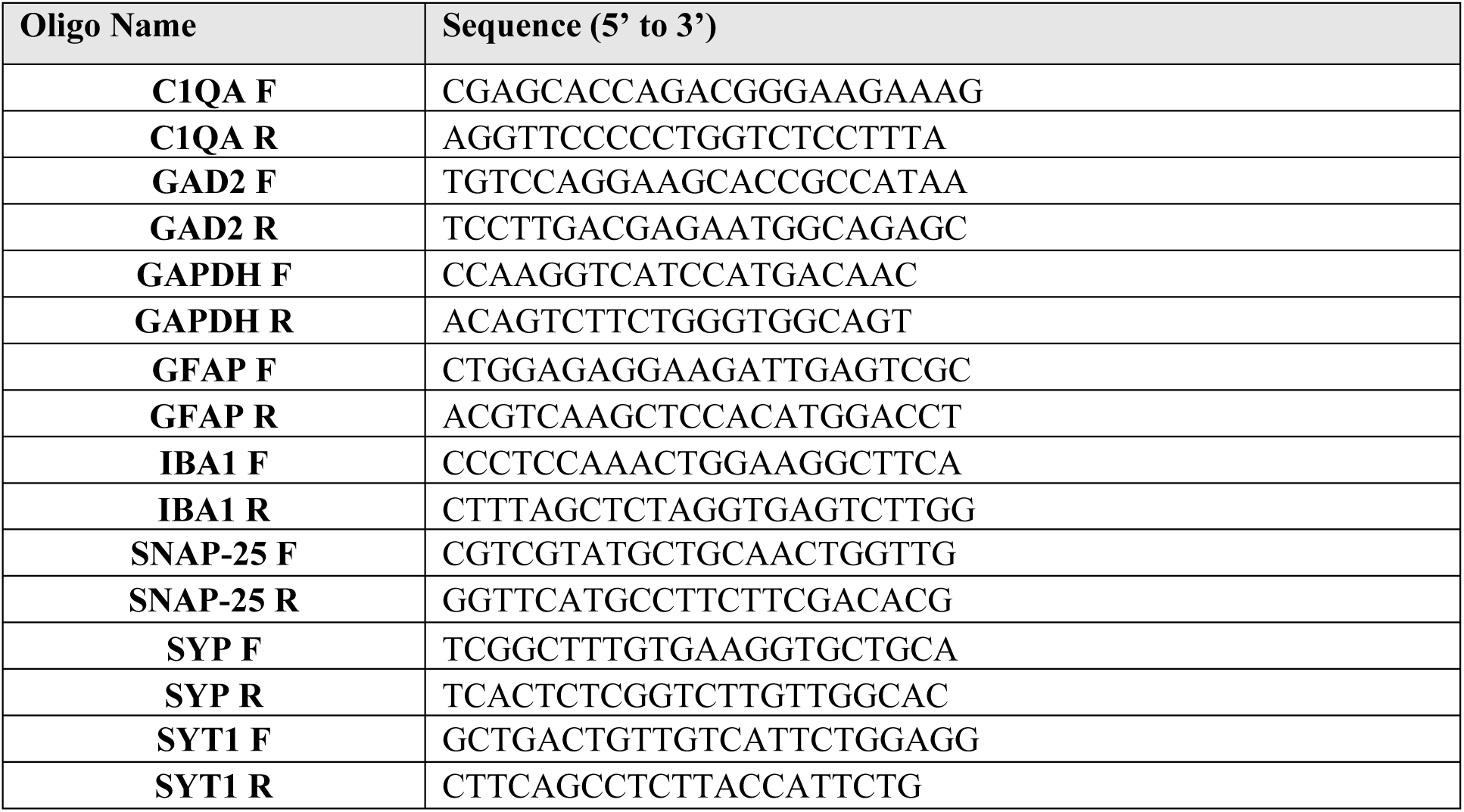
Primer sequences used for RT-qPCR analysis.

### Western blots

HBSCs were removed from the culture membrane with a scalpel blade into RIPA buffer (ThermoFisher Scientific: 89901) with protease inhibitor cocktail (1X) and EDTA (1X) (ThermoFisher Scientific: 78429). Slices were thoroughly homogenised via pipette trituration. A BCA assay was used to assess protein concentration, and normalised stocks were made. Stocks were then mixed into equal volumes of 2X Laemmli buffer (Merck: S3401-10VL) and boiled for 10 minutes at 98 °C. Each sample was loaded into 4-12% NuPage Bis-Tris gels (Invitrogen: NP0336BOX), before proteins were separated by electrophoresis using MES SDS running buffer (Invitrogen: NP0002). Proteins were then transferred onto PVDF transfer membranes using an iBlot machine (Invitrogen: IB24002). Following, a total protein stain (Li-Cor Biosciences: 926-11016) image was acquired using a Li-Cor Odyssey Fc machine and then de-stained. Membranes were subsequently blocked for 1 hour using PBS Intercept Blocking Buffer (Li-Cor Biosciences: 927-70001). Primary antibodies (see **Supplementary Table 5** for details) were diluted in PBS Intercept Blocking Buffer with 0.1% Tween-20 and incubated with membranes overnight at room temperature, with shaking. Membranes were washed three times for 5 minutes with PBS-Tween, then incubated in darkness for 2 hours with IRDye secondary antibodies (see **Supplementary Table 5** for details), specific to the corresponding primary antibody species. Membranes were washed 3x in PBS supplemented with 0.1%Tween, 1x in PBS and then imaged using a Li-Cor Odyssey Fc machine. Western blot images were analysed using Empiria Studio (Version 2.3), with the software generating background-subtracted intensity signals. Proteins of interest were normalised to housekeeping proteins run on the same membrane.

**Supplementary Table 5:** Antibodies used for western blots.

| Primary Antibody | Concentration | Source | Catalogue Number |
| --- | --- | --- | --- |
| Rabbit PSD-95 | 1:500 | Abcam | ab18258-100µl |
| Rabbit TUJ-1 | 1:2500 | Sigma | T2200 |
| Mouse Synaptophysin | 1:500 | Abcam | ab8049-500µl |
| Rabbit GAPDH | 1:2500 | Abcam | ab9485 |
| Rabbit PGP9.5 | 1:1000 | Abcam | ab108986-100µl |
| Rabbit YKL-40 | 1:1000 | Cell Signaling Technology | 47066S |
| Goat GFAP | 1:500 | Abcam | ab53554 |
| Rabbit Cyclophilin-B | 1:1000 | Abcam | ab16045 |
| Mouse APP | 1:500 | Merck Millipore | MAB348 |
| Secondary Antibody | Concentration | Source | Catalogue Number |
| IRDye 680RD Donkey Anti-Rabbit | 1:10000 | Li-Cor Biosciences | 926-68073 |
| IRDye 800CW Donkey Anti-Goat | 1:10000 | Li-Cor Biosciences | 926-32214 |
| IRDye 800CW Donkey Anti-Mouse | 1:10000 | Li-Cor Biosciences | 926-32212 |

### Statistics

All data was analysed using R and R Studio. Statistical tests were chosen according to the experimental design and dataset type.

The majority of statistical analyses used mixed effects models (‘lme4’ R package), as this allowed us to test if treatment group impacted our variable of interest while controlling for pseudoreplication by including random effects. The dataset was assessed for normality, linearity and homogeneity of variance. If the data failed these criteria, then the data was either transformed using the method that was best transformed each individual model to fit the assumptions or a generalised mixed effect model was used and the family for the model was changed to fit the distribution of the data. Data transformations included square root, log, arcsine square root, and Tukey transformation and data families included Gamma, Poisson and Binomial. *Post-hoc* testing was conducted for pairwise comparisons, estimated marginal means and 95% confidence intervals (’emmeans’ package), with Tukey correction for multiple comparisons. When data was transformed to meet the model assumptions then the reported effect sizes are computed on the back transformed data. Statistical details for each individual analyses can be found in the results text. Significance values were reported as *p* < 0.05 *, *p* < 0.01 *, *p* < 0.001 ***.

Correlation analyses were performed using the stats package. Datasets were assessed for normality and if these criteria were met, a Pearsons correlation was used. If not, a Spearmans correlation was used. Significance values are reported as *p* < 0.05 *, *p* < 0.01 *, *p* < 0.001 ***. Chi squared tests were performed using the stats package.

## Acknowledgements

We thank the Lothian NRS (NHS Research Scotland) BioResource and Tissue Governance unit and EMERGE Research Nurse team, particularly Allan Macraild, Ikeoluwa Adekoya, Anuka Boldbaatar and Sarah Risbridger, for obtaining informed consent and access cortical tissue samples from NHS patients. We thank the Alzheimer Scotland Brain and Tissue Bank and the Massachusetts Alzheimer’s Disease Research Centre (grant 1P30AG062421-01) for access to human post-mortem brain tissue. Thanks to the Genetics Core at the Edinburgh Clinical Research Facility, University of Edinburgh, for genotyping the human brain samples. We thank Dr Henner Koch, Dr Faye McLeod and Dr Daniel Erskine for their valuable discussions on protocol development and optimisation of human brain slice cultures. Most importantly, we would like to thank patients and their families for providing tissue donations, without which this work would not be possible. This work was funded by grants awarded to Dr Claire Durrant from Race Against Dementia (ARUK-RADF-2019a-001), The James Dyson Foundation, and the Alzheimer’s Society (581 (AS-PG-21-006), and by grants to Prof Tara Spires-Jones from the UK Demetia Research Institute, which receives its funding from DRI Ltd, funded by the UK Medical Research Council, Alzheimer’s Society, and Alzheimer’s Research UK (grant code UKDRI-Edin005). The confocal microscope was generously funded by Alzheimer’s Research UK (ARUK-EG2016A-6) and a Wellcome Trust Institutional Strategic Support Fund at the University of Edinburgh. Danilo Negro is funded by Wellcome Trust PhD programme in Translational Neurosciences (grant code: 228327/Z/23/Z). Dr Robert McGeachan is funded by the Wellcome Trust, as part of the Edinburgh Clinical Academic Track for Veterinary Surgeons (225442/Z/22/Z).

## Data availability

Data will be available on Edinburgh University Data Repository.

